# Characterization of intermolecular base pairing using AMT crosslinking in mammalian cells during oxidative stress

**DOI:** 10.1101/2025.05.21.655305

**Authors:** Jeffrey N. Liao, Isabel Betances, Emily Shen, Tatjana Trcek

## Abstract

Intermolecular base pairing is essential for normal gene regulation but can also cause neurodegeneration when dysregulated. Despite its importance, intermolecular base pairing remains poorly characterized in cells. We coupled AMT crosslinking with RNA pulldown to probe intermolecular base pairing in mammalian cells under oxidative stress, a state that drives G3BP1/2-dependent stress granule assembly, translational repression, and increased exposure of RNA sequences available for intermolecular interactions. AMT captured intermolecular base pairing interactions among reporter RNAs in vitro and the known interaction between the U3 snoRNA and pre-18S rRNA in cellulo. Although detection sensitivity was limited for endogenous mRNAs, we found that base pairing among inducibly expressed reporter mRNAs is sensitive to RNA sequence, oxidative stress, and G3BP1/2. Furthermore, chemical probing revealed that base accessibility to DMS within non-translated regions of candidate RNAs remains similar between stressed and unstressed cells, and those lacking G3BP1/2, indicating that RNAs remained state under these conditions. Our findings establish an approach for studying RNA base pairing in cells and suggest that intermolecular base pairing may act together with RNA structure to preserve RNAs during oxidative stress.

**Summary statement:** Intermolecular RNA:RNA base pairing is essential for gene regulation but can interfere with post-transcriptional regulation when dysregulated. Despite their importance, intermolecular base pairing interactions in cells have remained poorly characterized. Using AMT crosslinking, we show that intermolecular base pairing can be captured in mammalian cells during oxidative stress and is influenced by RNA sequence, stress state, and the stress granule proteins G3BP1/2. These findings establish a cellular approach for studying RNA base pairing in situ and suggest that cells regulate intermolecular base pairing interactions to preserve proper RNA function during stress recovery.

## Introduction

Intermolecular RNA:RNA base pairing is central to post-transcriptional gene regulation and RNA organization. For example, 22-nucleotide microRNAs pair with target mRNAs to downregulate them (O’Brien et al., 2018). Moreover, intermolecular base pairing-dependent dimerization of *oskar* mRNA enhances its enrichment to the *Drosophila* oocyte posterior, which is required for embryonic patterning (Jambor et al., 2011), while Human immunodeficiency virus-1 (HIV-1) replication depends on RNA base pairing (Dirac et al., 2002, Paillart et al., 1996, St Louis et al., 1998). However, unregulated base pairing can be harmful. Intermolecular interactions among trinucleotide-repeat-expansion RNAs associated with neurodegenerative disease can form nuclear foci (Jain and Vale, 2017) that inappropriately sequester RNA-binding proteins (Mankodi et al., 2001), and generate cytotoxic RNA aggregates (Das et al., 2023). Furthermore, introducing base pairing-competent sequences into germ granule mRNAs also impairs germ cell maintenance and embryogenesis in *Drosophila* (Tian et al., 2025). Cells must therefore monitor intermolecular base pairing, particularly under conditions that enhance it, to preserve normal gene expression.

One such condition is oxidative stress. At the cellular level, oxidative stress activates evolutionarily conserved signaling pathways that drive stress-specific gene expression responses, including modulation of protein synthesis. Specifically, it induces translational repression (Anderson and Kedersha, 2002, Aulas et al., 2017, Kedersha et al., 2002, Kedersha et al., 2020, Parker et al., 2022, Protter and Parker, 2016) which reduces global protein synthesis while permitting selective translation of stress-responsive genes (Pakos-Zebrucka et al., 2016). This increases the pool of ribosome-free (“naked”) mRNAs, which facilitates binding by two RNA-binding proteins (RBPs), RasGAP SH3-domain-binding protein 1 and 2 (G3BP1 and G3BP2; i.e. G3BP1/2) (Yang et al., 2020, Kedersha et al., 2020, Zhang et al., 2018). The interactions among G3BP1/2, and mRNAs promote the formation of extended intermolecular ribonucleoprotein (RNP) networks, driving assembly of large, mesoscale biomolecular condensates known as stress granules (SGs) (Bounedjah et al., 2014, Campos-Melo et al., 2021, Kedersha and Anderson, 2002, Kedersha et al., 2020, Panas et al., 2016, Ripin and Parker, 2023, Yang et al., 2020). These SGs help modulate mRNA translation during stress (Smith and Bartel, 2026) and promote cell survival (Fujikawa et al., 2023).

Translational repression also increases the potential for intermolecular RNA:RNA interactions, owing to both the spatial accumulation of ribosome-free mRNA and an increase in exposed sequences available for RNA:RNA interactions (Parker et al., 2025, Trussina et al., 2025, Van Treeck and Parker, 2018, Van Treeck et al., 2018). In support of this model, in vitro-transcribed RNAs, as well as cellular RNAs stripped of protein, can self-associate in the absence of protein (Aierken et al., 2024, Eisenberg and Felsenfeld, 1967, Jain and Vale, 2017, Langdon et al., 2018, Majumder et al., 2024, Parker et al., 2025, Tauber et al., 2020, Tian et al., 2025, Trussina et al., 2025, Van Treeck et al., 2018), demonstrating RNA’s intrinsic capacity for self-interaction. These interactions may involve both Watson-Crick base pairing and non-Watson-Crick mechanisms such as base stacking (Eisenberg and Felsenfeld, 1967, Van Treeck and Parker, 2018). While in vitro phase separation experiments show that RNAs can self-associate (Van Treeck and Parker, 2018, Van Treeck et al., 2018), imaging-based approaches suggest that intermolecular RNA:RNA interactions may also occur in cellulo during arsenite stress (Parker et al., 2025). However, enhanced RNA:RNA interaction leads to RNA entanglement and translation suppression in vitro (Trussina et al., 2025), suggesting that cells must tightly regulate such interactions to allow mRNAs to rejoin the translating pool once stress is relieved. While these studies demonstrate the presence of RNA:RNA interactions during stress, the formation of these interactions, specifically, Watson-Crick base pairing, has not been sufficiently characterized in cellulo. This raises the question of whether such interactions can be robustly detected and studied during oxidative stress in mammalian cells.

Chemical crosslinking is a widely used method for identifying RNA:RNA interactions, including base pairing, in cellulo. In particular, photochemical crosslinking with psoralen or psoralen derivatives such as 4′-aminomethyltrioxsalen (AMT) is commonly used because the crosslinks can be reversed (Velema and Lu, 2023), enabling downstream cDNA synthesis and analysis by qRT-PCR and next-generation sequencing.

In this study, we used AMT crosslinking followed by RNA affinity pulldown with biotinylated DNA oligonucleotides (Aw et al., 2016, Aw et al., 2017, Lu et al., 2018, Lu et al., 2016) to probe intermolecular base pairing in mammalian cells under sodium arsenite-induced oxidative stress. We found that AMT captures intermolecular base pairing interactions among reporter RNA fragments in vitro, in the absence of other cellular components, as well as the previously characterized intermolecular base pairing interaction between the U3 snoRNA and pre-18S rRNA in cellulo. Applying this approach to study interactions among mRNAs, we observed that, while the AMT crosslinking reaches the limits of its detection sensitivity when applied to endogenous mRNAs in cellulo, limiting our ability to study the regulation of intermolecular base pairing in finer detail, it nevertheless revealed that intermolecular base pairing among inducibly expressed reporter mRNAs is sensitive to RNA sequence, oxidative stress, and G3BP1/2 protein expression. Furthermore, applying chemical structure probing to monitor base accessibility, we found that base accessibility to DMS within the non-translating regions of *AHNAK* mRNA and lncRNA *NORAD*, two of the longest and most highly enriched stress granule transcripts, remains largely similar in stressed compared to unstressed cells, and in cells lacking G3BP1/2 expression. Together, our findings suggest that intermolecular base pairing in cells is regulated by RNA sequence, oxidative stress, and G3BP1/2, and that this regulation, together with the preservation of RNA structured state, may help maintain proper RNA function as cells recover from stress.

## Results

### AMT crosslinking coupled with RNA pulldown captures known pre-18S rRNA/U3 snoRNA base pairing interactions

To study intermolecular base pairing in cells, we utilized the psoralen derivative AMT for crosslinking, followed by affinity RNA pulldown with biotinylated DNA oligonucleotides, which was previously used to study base pairing interactions among mRNAs in human cells (Aw et al., 2016, Aw et al., 2017, Lu et al., 2018). Like psoralen, AMT crosslinks pyrimidines on adjacent RNA strands (Calvet and Pederson, 1979, Cimino et al., 1985, Nilsen, 2014, Stadler and Stafforst, 2014, Velema and Lu, 2023) (Fig. 1A), but it is more water-soluble (Nilsen, 2014) ensuring a higher effective concentration of the crosslinker in samples. Both, AMT and psoralen have been widely applied to map intermolecular RNA:RNA interactions in cells involving rRNAs, snRNAs, snoRNAs, mRNAs and long non-coding (lnc) RNAs (Aw et al., 2016, Lu et al., 2016, Sharma et al., 2016, Singh et al., 2024). While AMT cannot probe all possible intermolecular RNA:RNA interactions, we reasoned that investigating base pairing through AMT crosslinking nevertheless provides an insight into how cells may more broadly regulate intermolecular RNA:RNA interactions during oxidative stress.

**Figure 1:**
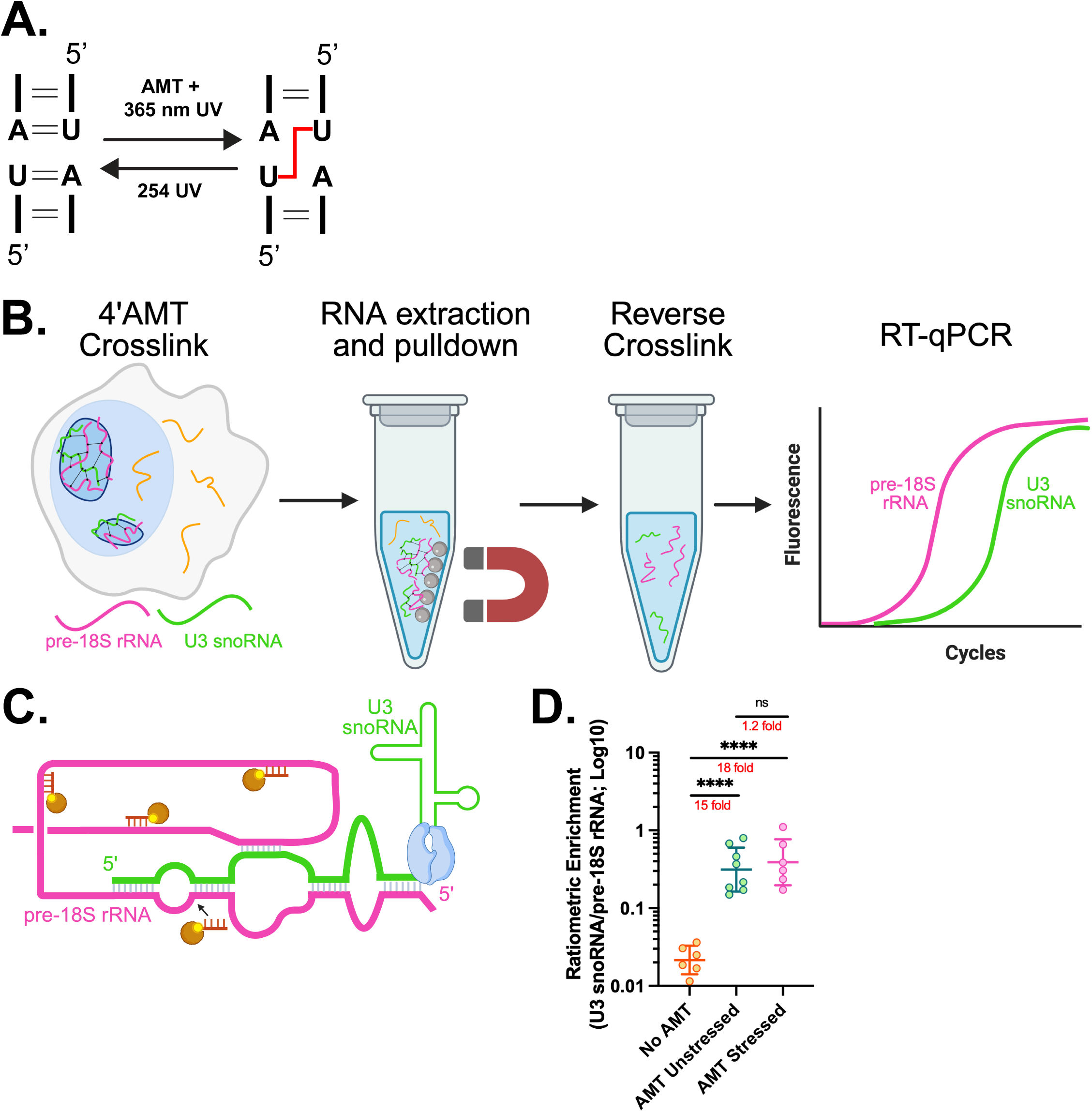
AMT crosslinking coupled with RNA pulldown captures known pre-18S rRNA/U3 snoRNA base pair interactions. **A.** Schematic of AMT crosslinking of pyrimidine dinucleotides adapted from Stadler et al (Stadler and Stafforst, 2014). **B.** Schematic of the AMT crosslinking and RNA pulldown approach using biotinylated oligonucleotides. **C.** Schematic of interactions between the nucleolar pre-18S rRNA and U3 snoRNA adapted from Sardana et al. (Sardana et al., 2015), with four biotinylated oligonucleotides + streptavidin beads (yellow/brown circles and lines) used to pulldown on the pre-18S rRNA shown. The levels of co-precipitated U3 snoRNA were quantified with RT-qPCR (blue polymerase). **D.** Ratiometric enrichment of U3 snoRNA levels normalized by the pre-18S rRNA levels in untreated, unstressed cells (“no AMT”) and in unstressed and stressed cells treated with AMT (“AMT Unstressed” and “AMT Stressed”, respectively). For no AMT and AMT-treated cells, a mean of six and eight independent samples +/-STDEV is plotted, respectively. Significance was determined using Welch’s ANOVA with Dunnett’s T3 multiple comparison correction. ****: p< 0.0001.

AMT crosslinking occurs upon exposure to 365 nm UV light, which is reversible with 254 nm UV light exposure (Fig. 1A) (Stadler and Stafforst, 2014). For affinity pulldown of intermolecular base pairing interactions, an AMT-crosslinked RNA is precipitated with biotinylated DNA oligos (Fig. 1B) (Torres et al., 2018, Aw et al., 2016). After reversal of crosslinking, the levels of co-precipitated RNA can be analyzed using quantitative reverse transcription PCR (RT-qPCR; Fig. 1B) (Aw et al., 2016), allowing for a quantitative evaluation of intermolecular base pairing.

To validate the feasibility of our crosslinking approach for detecting intermolecular base pairing in human osteosarcoma (U-2 OS) cells, we used the well-characterized pre-18S ribosomal RNA (rRNA)/U3 small nucleolar RNA (snoRNA) as a model system (Fig. 1C), whose intermolecular base pairing interactions occur in the nucleolus (Aw et al., 2016, Beltrame and Tollervey, 1992, Beltrame and Tollervey, 1995, Henras et al., 2015, Tschochner and Hurt, 2003). Psoralen crosslinking demonstrated that these two RNAs interact *via* regions of extended sequence complementarity (Fig. 1C) (Dutca et al., 2011, Maser and Calvet, 1989).

After reversing AMT crosslinks, we evaluated the levels of co-precipitated U3 snoRNA by RT-qPCR (Fig. 1B), normalizing against the background from non-specific RNA binding to magnetic beads during immunoprecipitation (Fig. S1A; Materials and Methods). To achieve this, we used biotinylated antisense oligos against GFP mRNA, which is not expressed in WT U-2 OS cells. This allowed us to evaluate non-specific co-precipitation, which can vary among experimental conditions (Materials and Methods), in agreement with previous observations (Aw et al., 2016). After this normalization, we calculated the ratio of the U3 snoRNA fold enrichment to the pre-18S rRNA fold enrichment (Fig. S1A), which allowed direct comparison of U3 snoRNA co-precipitation with pre-18S rRNA across experimental conditions (Materials and Methods). This approach revealed that, in unstressed cells, AMT crosslinking increased the ratiometric enrichment of U3 snoRNA with pre-18S rRNA 15-fold, a statistically significant increase (Fig. 1D, Fig. S1A). The efficiency of U3 snoRNA precipitation was evident from both the high ratiometric enrichment of U3 snoRNA relative to pre-18S rRNA (Fig. 1D) and the high fold enrichment of U3 snoRNA captured with U3 probes compared with GFP biotinylated probes (Fig. S1B,C). Importantly, the high levels of this enhancement persisted once arsenite stress was applied (Fig. 1D, Fig. S1B,C).

To assess the specificity of crosslinking and pulldown, we probed for the presence of the U2 small nuclear RNA (snRNA), a spliceosomal RNAs essential for mRNA splicing (Tholen, 2024, Tholen et al., 2022). Despite being expressed at levels ∼100-fold higher than U3 snoRNA, the U2 snRNA showed no significant co-precipitation with pre-18S rRNA in unstressed or stressed conditions (Fig. S1D-G).

Together, our experiments demonstrate the efficacy of the AMT approach in probing intermolecular base pairing within cells. Since interactions between the pre-18S rRNA and U3 snoRNA occur in the nucleolus (Henras et al., 2015, Tschochner and Hurt, 2003), our experiments further demonstrate that AMT crosslinking can capture intermolecular base pairing interactions within a biomolecular condensate in cellulo.

### AMT crosslinking enhances detection of intermolecular base pairing interactions between in vitro transcribed reporter RNAs

Next, we sought to investigate intermolecular base pairing of mRNAs in cells. Because the mRNAs that interact in cells are less understood, we engineered *Firefly* (*FLuc*) and *Renilla* (*RLuc*) luciferase reporter mRNAs that could report on such interactions. First, we tested AMT crosslinking-dependent detection of intermolecular base pairing between in vitro transcribed *FLuc* and *RLuc* (Fig. 2A,B). AMT crosslinking increased the co-precipitation of *RLuc* with *FLuc* coding sequences (CDS) 272-fold relative to the non-crosslinked control (Fig. 2C). This indicates that AMT enhanced detection of base pairing interactions between these two in vitro transcribed RNAs in the absence of other cellular components. Consistent with efficient crosslinking, both *FLuc* and *RLuc* RNA fragments contain many potential AMT-crosslinkable sites (Fig. S2A). Despite some variability in the Cq values for *FLuc* and *RLuc* co-precipitated with biotinylated GFP probes, the largest difference in Cq values was observed from pulldowns with the *FLuc* probes, which after normalization with the GFP probes, resulted in a noticeable change in fold enrichment (Fig. S2B,C). The increase in ratiometric enrichment therefore reflects enhanced AMT-dependent co-precipitation of the RNA, rather than an artifact of normalization to differences in background levels.

**Figure 2:**
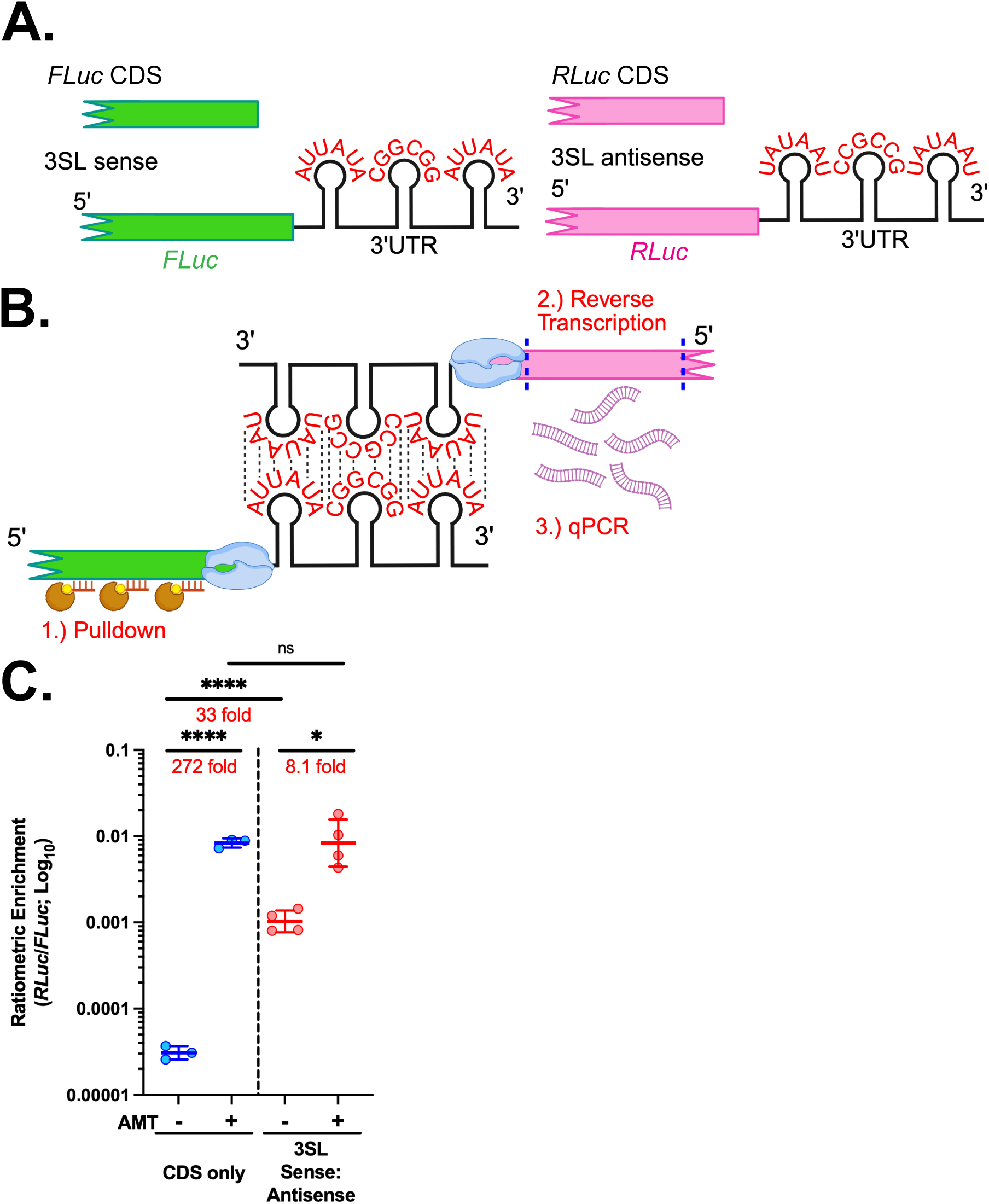
AMT crosslinking enhances detection of base pairing interactions of *in vitro* transcribed reporter RNAs. **A.** Schematic of the *RLuc* and *FLuc* luciferase reporters used for in vitro RNA precipitation. The green and magenta sections of the reporters represent portions of the *FLuc* and *RLuc* CDS regions, respectively. **B.** Schematic of interactions between the 3SL sense *FLuc* reporter and 3SL antisense reporter RNA, with three biotinylated oligonucleotides + streptavidin beads used to pulldown on the 3SL sense *FLuc* RNA. The levels of co-precipitated 3SL antisense *RLuc* RNA were quantified with RT-qPCR. **C.** Ratiometric enrichment of the *in vitro* transcribed *RLuc* CDS with *FLuc* CDS (blue), and *FLuc* 3SL sense with *RLuc* 3SL antisense (red) in the presence and absence of AMT. A mean of three (CDS) and four (CDS+3′UTR) independent samples +/- STDEV is plotted, respectively. Significance was determined using Welch’s ANOVA with Dunnett’s T3 multiple comparison correction. *: p < 0.05, ****: p < 0.0001.

Previous work from our lab and others has demonstrated that complementary sequences can promote intermolecular base pairing between both in vitro-transcribed reporter transcripts and endogenous mRNAs (Bose et al., 2024, Bou-Nader and Zhang, 2020, Clever et al., 1996, Ferrandon et al., 1997, Hussein et al., 2010, Jambor et al., 2011, Johnson and Telesnitsky, 2010, Langdon et al., 2018, Paillart et al., 2004, Tian et al., 2025, Tian and Trcek, 2026, Ye et al., 2026). Based on these observations and following the design we used to study intermolecular base pairing of *Drosophila* germ granule mRNAs (Tian et al., 2025), we modified the *FLuc* and *RLuc* reporter RNAs by fusing them to the 3′ UTR of the *Drosophila* gene *shutdown* (*shu*). This 3′ UTR also contained three stems derived from the SL1 motif, which is required for dimerization of human immunodeficiency virus (HIV) RNA (termed “3SL sense:antisense”; Fig. 2A) (Dirac et al., 2002, Paillart et al., 1996, St Louis et al., 1998). These stems contained AU- and GC-rich sequences that were both complementary between *FLuc* and *RLuc* and crosslinkable by AMT (Fig. 2B, Materials and Methods).

We evaluated the co-precipitation of in vitro transcribed 3SL sense:antisense RNA fragments, which contained the *shu* 3′ UTR and the last 705 and 281 nucleotides of the *RLuc* and *FLuc* CDSs, respectively. AMT crosslinking increased the co-precipitation of *RLuc* with *FLuc* 3SL sense:antisense fragments 8.1-fold relative to the non-crosslinked control, a statistically significant increase (Fig. 2C). The ratiometric enrichment in the presence of AMT was similar to that of the CDS-only fragments.

In the absence of AMT, however, the enrichment of *RLuc* with *FLuc* 3SL sense:antisense was nearly 33-fold higher than for the CDS only fragments (Fig. 2C). While we observed small differences in Cq values of co-precipitated *RLuc* and *FLuc* with biotinylated GFP probes with and without AMT, we also observed a large difference in Cq value for co-precipitated *RLuc* in the absence of AMT for the 3SL sense:antisense reporter (Fig. S2D). This resulted in a noticeable increase in the *RLuc* fold enrichment (Fig. S2E) and subsequently higher ratiometric enrichment (Fig. 2C). This is consistent with more stable base pairing that does not require crosslinking.

Together, these results indicate that intermolecular base pairing of in vitro transcribed RNAs can occur in the absence of AMT-mediated crosslinking. However, AMT and RNA sequence strongly enhance the detection of these interactions, enabling efficient co-precipitation of interacting RNA molecules.

### G3BP1/2, MCP-G3BP1 expression, and oxidative stress modulate AMT-dependent detection of base pairing interactions among reporter mRNAs in cells

Next, we investigated intermolecular interactions between the luciferase reporters in unstressed and stressed cells. Because the 3SL sense:antisense reporter RNAs showed a higher baseline of co-precipitation even without AMT, suggesting more stable base pairing (Fig. 2C, Fig. S2D,E), we examined co-precipitation of this reporter in cells. We also inserted twelve bacteriophage MS2 stem-loops at the end of the 3′ UTRs of both reporter mRNAs (Fig. 3A) to recruit proteins fused to the MS2 coat protein (MCP) to the mRNAs (see below). We then expressed these reporter RNAs under tetracycline (tet)-inducible promoters in tet-on (i.e. EF1αTET) U-2 OS cells. Single-molecule fluorescence in situ hybridization (smFISH) confirmed co-expression of both reporters (Fig. 3B). We also confirmed that AMT and UV crosslinking did not disrupt SGs (Fig. S3A), indicating that these treatments did not grossly interfere with cellular biology.

**Figure 3:**
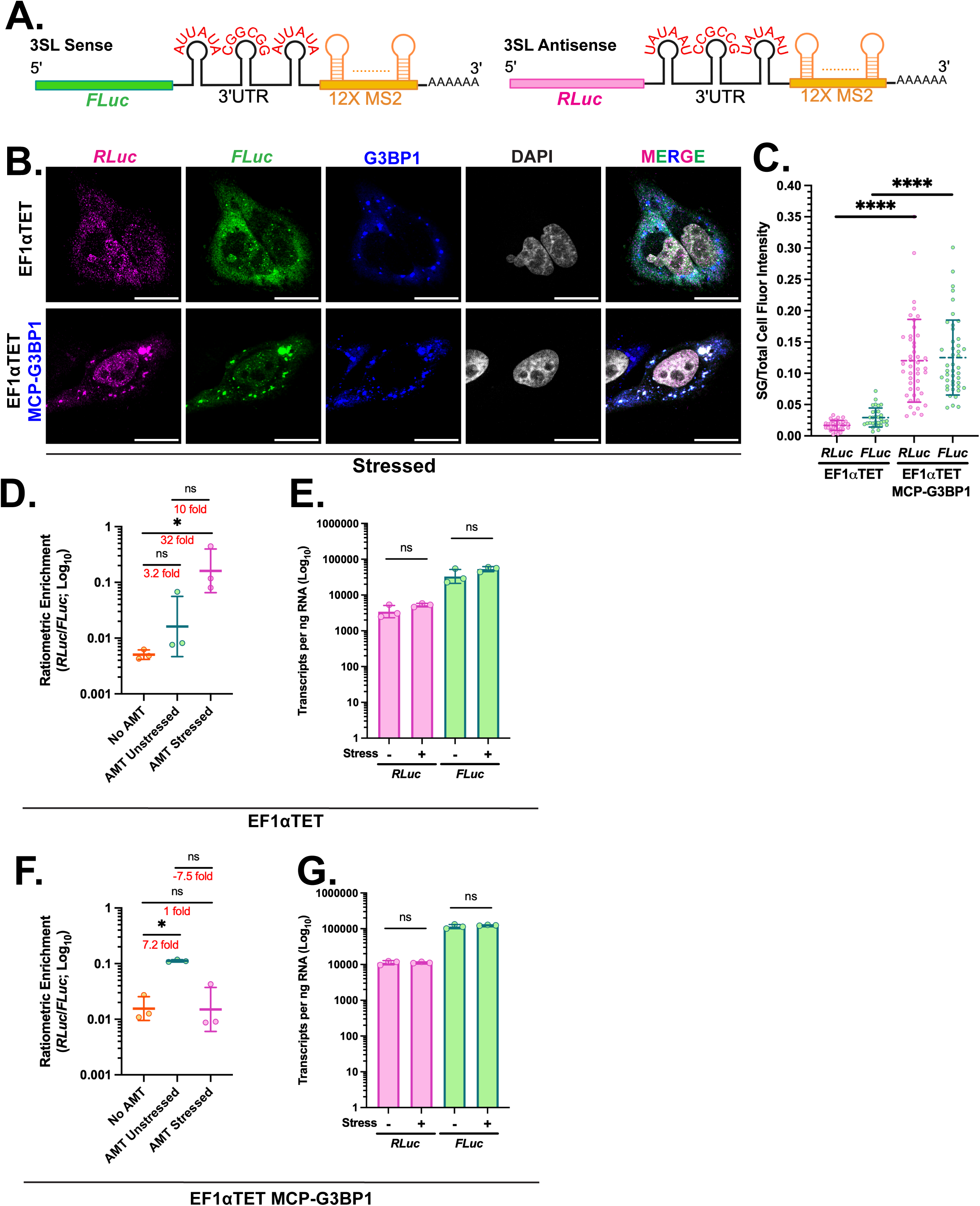
Oxidative stress and MCP-G3BP1 co-expression modulate AMT-dependent detection of base pairing interactions among reporter mRNAs in cellulo. A. Schematic of the 3SL sense:antisense *RLuc* and *FLuc* reporter mRNAs tagged with 12 MS2 stem loops in their 3′UTRs. B. Images of cells immunostained for G3BP1 protein (blue) and smFISH-stained *RLuc* (magenta) and *FLuc* (green) mRNAs in stressed EF1αTET or EF1αTET MCP-G3BP1 U-2 OS cells. The antibody recognizes both endogenous G3BP1 and ectopically expressed 6xHis-MCP-G3BP1. DAPI stain demarcates the nuclei (white). Scale bar: 20 µm. C. Fraction of *FLuc* and *RLuc* mRNA signals localized within SGs relative to total cellular signal in WT (green) and 6xHis-MCP-G3BP1-expressing (magenta) cells. A total of 31 WT and 44 MCP–G3BP1–expressing cells were analyzed with the mean per condition ± SD plotted. Statistical significance was examined using Student’s T-test with Welch’s correction. D. Ratiometric enrichment of the co-precipitated 3SL sense:antisense *RLuc* and *FLuc* reporter pair in unstressed EF1αTET cells treated with PBS and in unstressed and stressed EF1αTET cells treated with AMT. Significance was determined using Welch’s ANOVA with Dunnett’s T3 multiple comparisons. Mean of three biological replicates +/- STDEV is plotted. E. RT-qPCR analysis of *FLuc* and *RLuc* mRNA levels after 36 hours of 1ug/mL doxycycline induction in stressed and unstressed EF1αTET cells. The mean of three biological replicates ± STDEV is shown for log transformed values. qPCR quantification was done using absolute standard curves generated from dilutions of purified DNA amplicons. Significance was determined using Student’s T-test with Welch’s correction. F. Ratiometric enrichment of the co-precipitated 3SL sense:antisense *RLuc* and *FLuc* reporter pair in unstressed MCP-G3BP1 expressing EF1αTET cells treated with PBS and in unstressed and stressed MCP-G3BP1 expressing EF1αTET cells treated with AMT. Mean of three biological replicates +/- STDEV is plotted. **G.** RT-qPCR analysis of *RLuc* and *FLuc* mRNA levels after 36 hours of 1ug/mL doxycycline induction in stressed and unstressed EF1αTET MCP-G3BP1 cells. ns: not significant, *: p < 0.05, ****: p < 0.0001.

AMT treatment did not significantly enhance the co-precipitation of 3SL sense:antisense *RLuc* with *FLuc* in unstressed WT cells (Fig. 3D). However, after treatment with sodium arsenite, we observed a statistically significant 32-fold increase in co-precipitation in stressed cells treated with AMT, compared with unstressed cells not treated with AMT (Fig. 3D). This increase was not explained by changes in reporter abundance, as the levels of 3SL sense:antisense *RLuc* and *FLuc* mRNAs were largely similar in stressed and unstressed cells (Fig. 3E). Our data are consistent with enhanced AMT-dependent co-precipitation of 3SL sense:antisense *RLuc* with *FLuc* during stress.

However, the Cq value of *RLuc* co-precipitated with biotinylated FLuc probes during stress was at most 1.5 cycles lower than with biotinylated GFP probes (Fig. S3B), corresponding to only a small, ∼ 2.8 fold enrichment (Fig. S3C). Therefore, although AMT-dependent capture of base pairing interactions between 3SL sense:antisense *RLuc* and *FLuc* reporter mRNAs was statistically significant, the detection of these interactions began to approach the sensitivity limit of our approach.

Furthermore, co-precipitation of 3SL sense:antisense *RLuc* with *FLuc* did not significantly increase during oxidative stress in EF1αTET cells lacking G3BP1/2 (i.e. ΔΔG3BP1/2 cells), as reflected in both the ratiometric enrichment and the fold enrichment of *RLuc* with *FLuc* mRNAs (Fig. S3D-G). We confirmed the lack of G3BP1/2 expression in ΔΔG3BP1/2 cells and that these cells failed to form SG-like structures (Fig. S4A,B), consistent with previous observations (Yang et al., 2020). Reporter expression was comparable in ΔΔG3BP1/2 cells before and after oxidative stress (Fig. S3E), indicating that reporter mRNA levels did not change in response to stress in these cells. Together, these results suggest that the stress-dependent increase in AMT-dependent detection of interactions between 3SL sense:antisense reporter mRNAs, which in EF1αTET U-2 OS cells rose above the detection limit of our assay, requires G3BP1/2, and possibly SG formation. Finally, we co-expressed the *RLuc* and *FLuc* reporters with MCP-G3BP1, following established MS2-mRNA-MCP-G3BP1 tethering protocols (Fischer et al., 2020). We confirmed that MCP-G3BP1 formed SGs and recruited canonical SG components (Fig. S4C-F; Materials and Methods), indicating that it retained key properties of endogenous G3BP1. In arsenite-treated cells, MCP-G3BP1 expression increased the enrichment of *RLuc* and *FLuc* reporter mRNAs in SGs, consistent with MCP-mediated recruitment of G3BP1 to the MS2-tagged mRNAs (Fig. 3B,C).

Under these conditions, AMT treatment of unstressed cells increased the co-precipitation of 3SL sense:antisense *RLuc* with *FLuc* 7.2-fold compared with unstressed cells not treated with AMT, a statistically significant change (Fig. 3F; Fig. S4G,H). Following oxidative stress, however, *RLuc* co-precipitation decreased to levels similar to those in unstressed cells without AMT (Fig. 3F, Fig. S4G,H). This decrease was not attributable to reduced reporter expression, as *FLuc* and *RLuc* mRNA levels remained unchanged after arsenite treatment (Fig. 3G). It is also unlikely to reflect a general impairment of AMT’s ability to crosslink RNAs during oxidative stress, as we co-precipitated the reporters in stressed WT cells (Fig. 3D) and AMT-dependent co-precipitation of U3 snoRNA with pre-18S rRNA persisted after arsenite treatment (Fig. 1D).

Together, our results suggest that in cells, AMT-dependent detection of intermolecular base pairing between MS2-tagged 3SL sense:antisense *RLuc* and *FLuc* reporter mRNAs is influenced by oxidative stress, by G3BP1/2, and by MCP-G3BP1 co-expression. In WT cells and in the absence of MCP-G3BP1, oxidative stress enhanced the detection of these interactions, and this enhancement required G3BP1/2. In contrast, co-expression of MCP-G3BP1 enhanced their detection already in unstressed cells, whereas oxidative stress then diminished, rather than further increased, reporter RNA co-precipitation.

### RNA sequence modulates AMT-dependent detection of base pairing interactions among reporter mRNAs in cells

Finally, we observed that the stress-dependent enhancement of *RLuc* co-precipitation with *FLuc* in WT cells was specific to the 3SL sense:antisense reporter pair. The *shu FLuc* and *RLuc* reporter pair, which lacked the stems containing complementary AU- and GC-rich sequences (Fig. S5A), did not show detectable co-precipitation under any condition. Instead, its co-precipitation levels were indistinguishable from those obtained with biotinylated GFP probes (Fig. S5B–E).

The MCP-G3BP1-dependent increase in co-precipitation that we observed in unstressed cells (Fig. 3F) was also specific to the 3SL sense:antisense reporters. No comparable statistically significant increases in co-precipitation were observed for *shu RLuc* with *FLuc* in unstressed or stressed MCP-G3BP1-expressing cells, compared with unstressed cells not treated with AMT (Fig. S6A–F). In arsenite-treated cells, MCP-G3BP1 expression increased the enrichment of *shu RLuc* and *FLuc* reporter mRNAs in SGs, consistent with MCP-mediated recruitment of G3BP1 to the MS2-tagged mRNAs (Fig. S6A,B). The levels of *shu RLuc* and *FLuc* mRNAs did not change across cellular conditions and cell types (Fig. S6F), and were similar to those of the 3SL sense:antisense reporters under the same conditions (Fig. 3G). The lack of AMT-dependent co-precipitation between *shu RLuc* and *FLuc* therefore cannot be explained by differences in expression levels.

Thus, in contrast to the 3SL sense:antisense reporters, *shu RLuc* and *FLuc* did not generate co-precipitation above the detection limit of our approach. This also contrasts with our in vitro results, where AMT crosslinking enhanced the co-precipitation of CDS-only fragments lacking the 3SL stems to levels similar to those of fragments containing the stems (Fig. 2C). These data suggest that AMT-dependent detection of intermolecular base pairing in cells is more stringent and depends, at least in part, on RNA sequence composition.

### AMT crosslinking has limited ability to detect intermolecular base pairing between the lncRNA *NORAD* and *AHNAK* mRNA during oxidative stress

We next applied AMT crosslinking to examine co-precipitation of endogenous mRNAs in cells before and during oxidative stress. To this end, we probed co-precipitation between *AHNAK* mRNA and the long non-coding (lnc) RNA *NORAD* as a representative RNA pair that may engage in intermolecular base pairing during oxidative stress. Both are among the longest cellular transcripts as well as among the most enriched in SGs (Khong et al., 2017), features thought to increase RNA’s propensity for intermolecular base pairing (Van Treeck and Parker, 2018, Van Treeck et al., 2018). In addition, thermodynamic predictions suggested that *NORAD* contains two regions located at its 5′ and 3′ ends capable of intermolecular base pairing with multiple regions across the *AHNAK* transcript (Fig. S7A; Materials and Methods). Moreover, both *AHNAK* and *NORAD* were replete with pyrimidine dinucleotides (Fig. S7B,C), which are typical sequence substrates for AMT crosslinking (Calvet and Pederson, 1979, Cimino et al., 1985, Nilsen, 2014, Stadler and Stafforst, 2014, Velema and Lu, 2023). Together, these features made the two transcripts well suited for probing AMT-dependent intermolecular base-pairing interactions in cellulo.

Because the specific regions involved in the potential *AHNAK–NORAD* interactions in cellulo were unknown, we designed a pool of 16 biotinylated DNA oligonucleotides spanning the entire *AHNAK* mRNA sequence (Fig. S7D, Table S1, Materials and Methods). To maximize the likelihood of identifying regions of *NORAD* that may base-pair with *AHNAK*, we designed 11 RT-qPCR amplicons spanning the entire *NORAD* sequence (Table S2). Characterization of the corresponding primers confirmed that the majority of amplicons were amplified with comparable efficiency (Fig. S7E).

We detected statistically significant increases in AMT-dependent ratiometric enrichment of a subset of *NORAD* regions with *AHNAK* in unstressed and stressed WT U-2 OS cells, compared with unstressed cells not treated with AMT (Fig. S7F). The *NORAD* regions detected in the co-precipitate, specifically amplicons 1, 2 and 8, were close to regions predicted to interact by thermodynamic modeling (Fig. S7A), consistent with the possibility that these regions contribute to intermolecular base pairing detectable by AMT crosslinking. However, the fold enrichment of *NORAD* in co-precipitates obtained with biotinylated *AHNAK* probes, relative to those obtained with biotinylated GFP probes, was close to baseline for all *NORAD* regions (Fig. S7G). We observed the same trend for co-precipitation of these two RNAs in ΔΔG3BP1/2 cells (Fig. S7H,I). In these cases, the ratiometric enrichment was more heavily influenced by the pulldown efficiency of *AHNAK*, rather than by co-precipitation of *NORAD*. Therefore, these results suggest that the abundance of intermolecular base pairing interactions between *NORAD* and *AHNAK* may be too low to reliably conclude whether these two transcripts base pair with each other in unstressed or stressed cells (see Discussion).

### *AHNAK* mRNA and lnc *NORAD* show a modest increase in spatial overlap during oxidative stress

The limited sensitivity of AMT crosslinking for detecting intermolecular base pairing between *NORAD* and *AHNAK* prevented us from examining how cellular stress modulates interactions between these endogenous RNAs. To circumvent this limitation, we instead examined the spatial organization of *NORAD* and *AHNAK* in stressed and unstressed cells using single-molecule fluorescence in situ hybridization (smFISH). Although smFISH alone cannot report on intermolecular interactions, it measures co-localization between molecules, allowing us to assess how RNAs are positioned relative to each other within subcellular environments, as we have done before (Trcek et al., 2020, Trcek et al., 2015, Tian et al., 2025). To define a reference for the spatial overlap expected between two RNAs in close proximity, we measured the co-localization (Pearson correlation coefficient, r) of two fluorescent signals originating from the same, doubly labeled *AHNAK* mRNA (Fig. 4A, B). This was r = 0.67 ± 0.06 (Fig. 4B, C). This value was almost identical to the one we recently reported for *oskar* mRNA, which engages in intermolecular base pairing interactions within *oskar* mRNA clusters in *Drosophila* (Ye et al., 2026). We therefore used the co-localization of doubly labeled *AHNAK* as a reference for the overlap expected between two RNAs brought into close proximity through intermolecular interactions. Because both signals come from a single molecule, this reference is likely close to the upper limit for two distinct RNAs.

**Figure 4:**
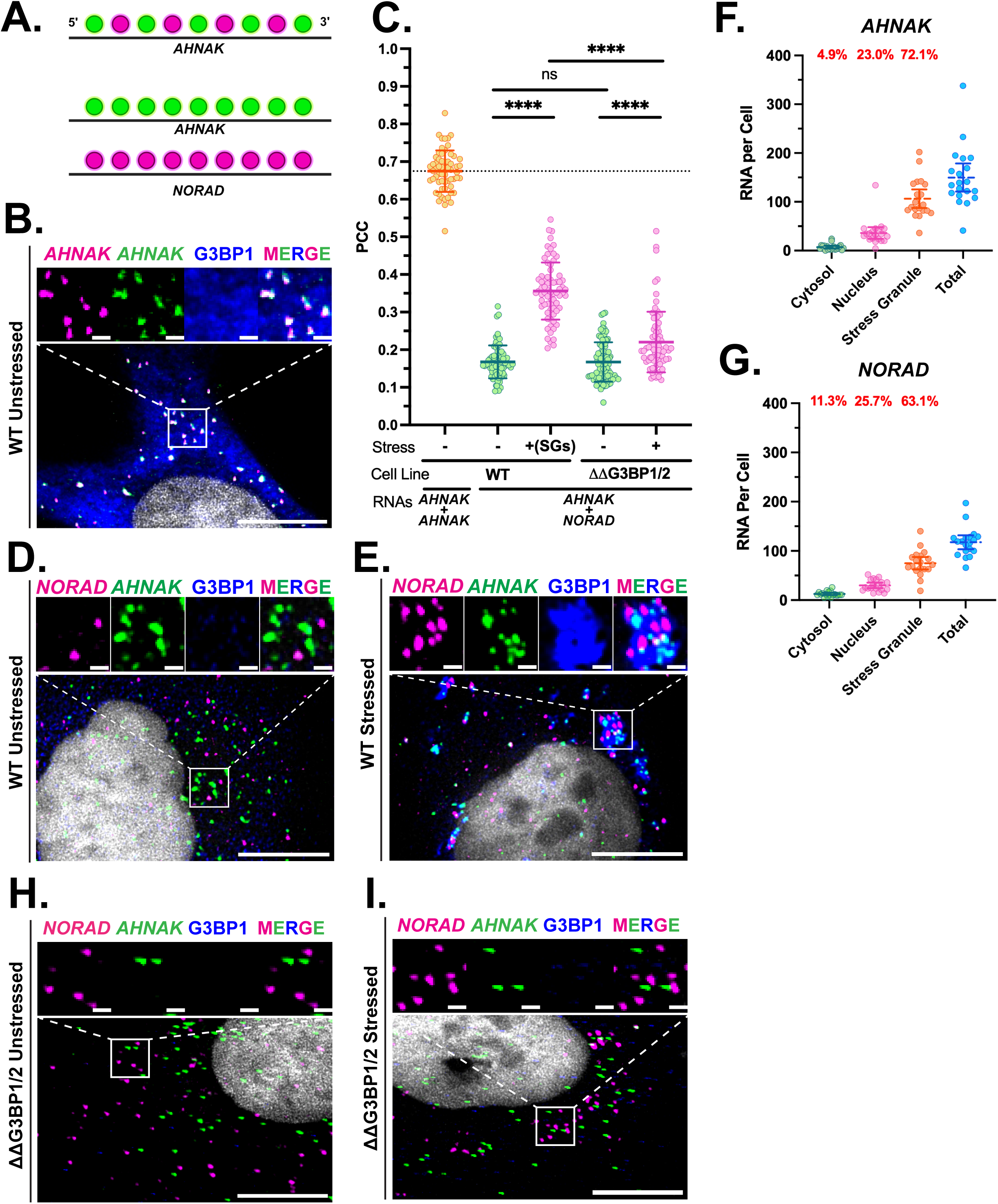
*AHNAK* mRNA and lnc *NORAD* show a modest increase in spatial overlap during oxidative stress. **A.** Schematic of fluorescent smFISH probes used to label *AHNAK* and *NORAD* RNAs. The dual labeled *AHNAK* mRNA consist of two probes sets interspersed between one another. **B.** Image of a doubly-stained *AHNAK* mRNA (magenta and green) with immunostaining for G3BP1 protein (blue) in unstressed WT cells. DAPI stain demarcates the nuclei (white). Scale bars: 1 and 10 µm in small and big images, respectively. Top: zoomed-in image of the white square shows co-localization of the fluorescent signal of the doubly-stained *AHNAK* mRNA in WT cells. **C.** Co-localization of doubly-stained *AHNAK* (orange circles) in WT unstressed cells and of *AHNAK* with *NORAD* in the cytoplasm of unstressed (green circles) and stressed (magenta circles) WT and ΔΔG3BP1/2 U-2 OS cells. For each condition, minimally 60 cells were analyzed. Statistical significance was examined using Welch’s ANOVA with Dunnett’s T3 multiple comparisons correction. ****: p < 0.0001. **D,E.** Images of cells smFISH-stained *NORAD* lncRNA (magenta) and *AHNAK* mRNA (green) in unstressed (D) and stressed (E) WT U-2 OS cells with immunostained for G3BP1 protein (blue). DAPI stain demarcates the nuclei (white). Zoomed-in images (white squares) show *AHNAK* and *NORAD* dispersed in the cytosol (D) or co-localized in SGs (E). **F, G.** Quantification of smFISH-stained *AHNAK* (F) and *NORAD* (G) in different cellular compartments in stressed WT U-2 OS cells. In stressed conditions, the cytoplasmic compartment represents the cytosol, excluding SGs. Percent of *AHNAK* (F) or *NORAD* (G) per compartment is shown above the graph. **H,I.** Images of cells immunostained for G3BP1 protein (blue) and smFISH-stained *AHNAK* (green) and *NORAD* (magenta) in unstressed (H) and stressed (I) ΔΔG3BP1/2 U-2 OS cells. DAPI stain demarcates the nuclei (white). Zoomed-in images in H and I (white squares) show *AHNAK* and *NORAD* dispersed in the cytosol. Scale bars: 1 and 10 µm in small and big images, respectively.

Our analysis revealed that 72.1% of *AHNAK* and 63.1% of *NORAD* localized to SGs in stressed WT U-2 OS cells (Fig. 4E-G), consistent with previous measurements (Khong et al., 2017). We also observed an increase in their co-localization within SGs, from r = 0.17±0.04 in the cytosol of unstressed cells to r = 0.36±0.07 in SGs (Fig. 4C). Although this increase was statistically significant, it was much lower than the co-localization coefficient recorded for doubly labeled *AHNAK* mRNA (Fig. 4C). Moreover, the corresponding *R*² values indicated that the enrichment into SGs explained less than 13% of spatial variability of *AHNAK* localization with *NORAD* (R^2^ = 0.36^2^*100%). Thus, although *AHNAK* and *NORAD* showed increased co-localization during stress, the extent of their spatial overlap remained modest relative to the reference.

Moreover, both transcripts remained mostly dispersed in stressed ΔΔG3BP1/2 U-2 OS cells without a detectable granule-like localization pattern (Fig. 4H,I). Although we recorded a statistically significant increase in co-localization between the two RNAs in stressed compared to unstressed ΔΔG3BP1/2 U-2 OS cells (r = 0.16±0.05 versus r = 0.22±0.08, respectively; Fig. 4C), the corresponding R² value indicated that co-localization in SGs accounted for less than 5% of the spatial variability in *AHNAK* localization with *NORAD* (R^2^ = 0.22^2^*100%). We observed a similar modest co-localization between *AHNAK* and two additional SG-enriched mRNAs, *TLN1* and *DYNC1H1*, in stressed WT and ΔΔG3BP1/2 U-2 OS cells (Fig. S8A-L). Both *TLN1* and *DYNC1H1* are among the longest and most highly enriched mRNAs in SGs (Fig. S8E,F) (Khong et al., 2017). Importantly, the modest co-localization was not attributable to reduced RNA expression in stressed cells, as levels of *AHNAK*, *NORAD*, *TLN1*, and *DYNC1H1* remained unchanged following arsenite treatment (Fig. S8M).

Together, these imaging experiments indicate that co-localization of *AHNAK* with *NORAD*, *TLN1*, and *DYNC1H1* increases modestly during stress, and that G3BP1/2 are required for these RNAs to localize to granules. The spatial overlap between these transcripts nevertheless remains modest during oxidative stress, even in stressed cells where they are recruited to SGs, an observation that does not support extensive intermolecular contacts between these RNAs in cells.

### DMS-MaPseq probing revealed modest changes in base accessibility of the *AHNAK* 3′UTR during oxidative stress

Previous work from our lab showed that the abundance of intermolecular base pairing among mRNAs enriched in *Drosophila* germ granules, another biomolecular condensate characterized by high local RNA concentration, is constrained by RNA structure (Tian et al., 2025). Therefore, we speculated that RNA structure could play a role in limiting intermolecular base pairing during oxidative stress in mammalian cells.

To investigate this possibility, we used dimethyl sulfate mutational profiling followed by sequencing (DMS-MaPseq) to examine the accessibility of RNAs to chemical probing, as we have done in *Drosophila* (Tian et al., 2025). DMS chemically methylates single, unpaired adenines and cytosines (Tijerina et al., 2007, Lawley and Brookes, 1963) generating mutations during reverse transcription that can be detected using next-generation sequencing (Rouskin et al., 2014, Zubradt et al., 2017) and used to infer secondary and tertiary RNA structure (Tomezsko et al., 2020, Rouskin et al., 2014, Lange et al., 2024). However, in this study, we focused on examining changes in overall base accessibility to DMS to account for potential factors other than base pairing, such as protein binding and molecular crowding.

As a control, we first mapped the reactivity profile of the 5.8S ribosomal rRNA (5.8S rRNA), a non-coding RNA component of the large ribosomal subunit (AbouElela and Nazar, 1997) in unstressed U-2 OS cells. After normalizing the DMS signal (Materials and Methods), we observed a strong DMS signature on the RNA (Fig. 5A). Using these data, we overlaid the DMS mismatch ratio of bases >0.01 onto the previously recorded structure of the 5.8S rRNA (Fig. 5B) (The et al., 2017). We observed that A and C bases that reacted with DMS matched those that we previously identified as unpaired within the 5.8S rRNA (Fig. 5B). This result, together with the fact that DMS efficiently recapitulates the structure of the yeast 18S rRNA to 94% accuracy *in vivo* (Rouskin et al., 2014), demonstrated the effectiveness of the DMS-MaPseq in probing the base accessibility of RNAs within a proteinaceous environment.

**Figure 5:**
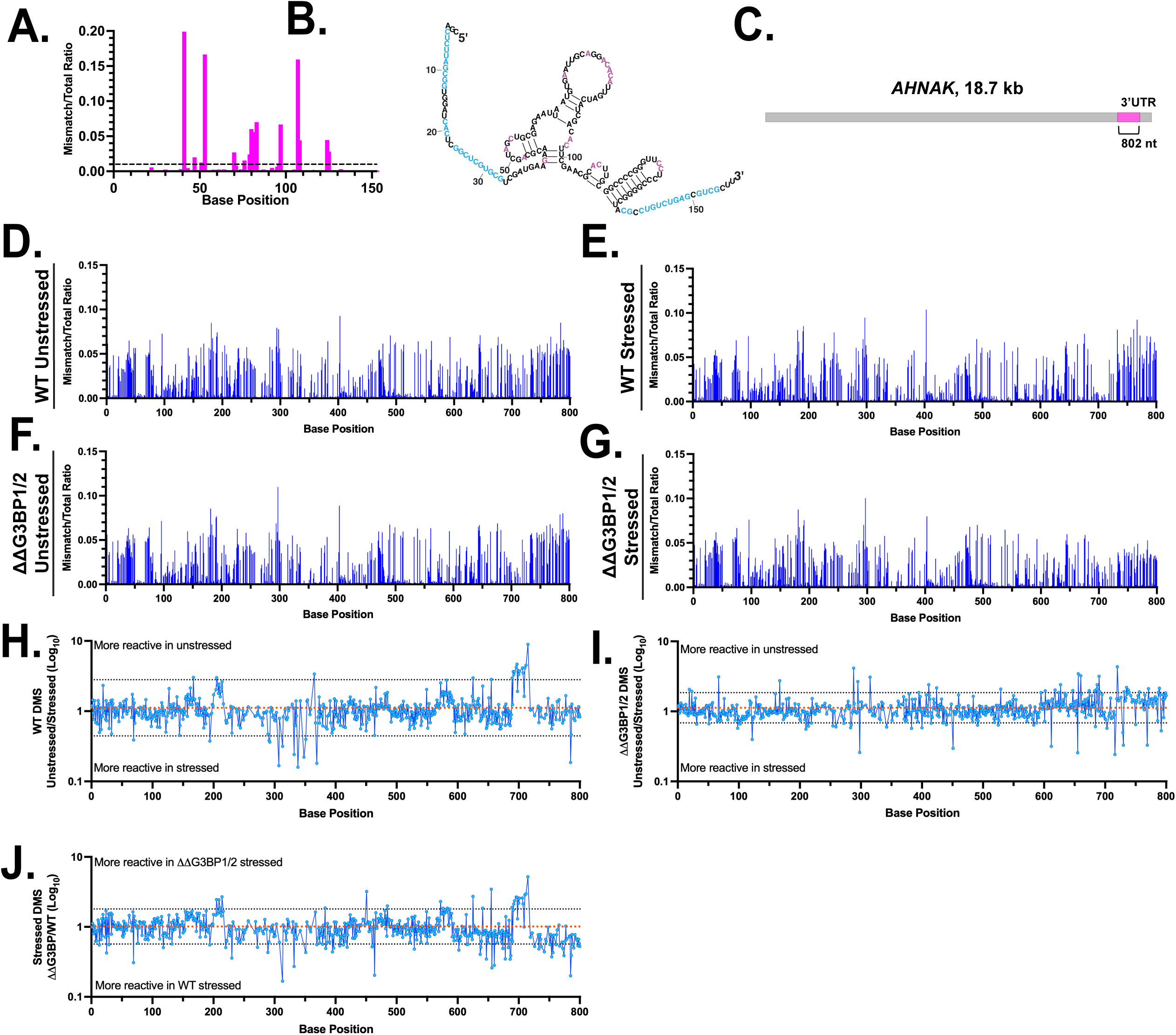
DMS-MaPseq probing revealed modest changes in base accessibility of the *AHNAK* 3′UTR during oxidative stress. **A.** Per base ratio between the number of reads containing a mismatch versus total number of reads revealed by the Induro reverse transcriptase for the human 5.8S rRNA sequence. The dotted line demarcates a mismatch ratio >0.01. **B.** Predicted secondary structure of the 5.8S rRNA generated from the probe accessibility data obtained in Fig. 5A. Bases marked in blue are base paired with the 28S rRNA (The et al., 2017). Bases marked in pink display a mismatch ratio >0.01 (from Fig. 5A). **C.** Schematic of *AHNAK* mRNA 3′UTR region (magenta) assayed with DMS-MaPseq. **D-G.** DMS mismatch ratio of the *AHNAK* 3′UTR in unstressed (D) and stressed WT cells (E) and unstressed (F) and stressed (G) ΔΔG3BP1/2 cells. The mismatch ratio represents ratio of reads with DMS modification-induced mismatches at a particular base when compared to the reference transcriptome. The ratio shown is an average of three biological replicates. **H-J.** DMS ratios in the average per base DMS reactivity of the *AHNAK* 3′UTR in unstressed and stressed WT U-2 OS cells (H), unstressed and stressed ΔΔG3BP1/2 U-2 OS cells (I) and stressed ΔΔG3BP1/2 and stressed WT U-2 OS cells (J). The orange dotted lines denote the average ratio between two compared conditions while the black dotted lines represent the statistical significance threshold where bases that fall above and below the line show significant differences between the conditions examined. The ratio is calculated from the averages of the three biological replicates per condition (see Materials and Methods).

Next, we mapped the reactivity profile of *AHNAK* mRNA. Since the release of ribosomes from mRNAs during oxidative stress (Anderson and Kedersha, 2002, Aulas et al., 2017, Kedersha et al., 2002, Kedersha et al., 2020, Parker et al., 2022, Protter and Parker, 2016) is accompanied by increase in RNA double strandedness and changes in RNA structure (Qu et al., 2011, Amiri and Noller, 2019, Xie and Chen, 2017, Farenhem et al., 2025, Wu et al., 2024, Zhu et al., 2025) we specifically focused on mapping 802 of the 842 nucleotides (nts) of the *AHNAK* 3′UTR (Fig. 5C).

To separate the effects of oxidative stress from those of SGs on base accessibility, we compared individual biological replicates obtained from unstressed and stressed ΔΔG3BP1/2 cells to each other. We observed a high correlation of the average base accessibility to DMS for *AHNAK* 3′UTR among replicates, ranging from 0.97 to 0.99 in unstressed and from 0.96 to 0.98 in stressed cells (Fig. S9A,B). This indicated high reproducibility among biological replicates for each cellular condition.

Our measurements revealed similar absolute levels of DMS reactivity across conditions (Fig. 5D-G). Looking at the fold change in DMS reactivity of the *AHNAK* 3′UTR between unstressed and stressed ΔΔG3BP1/2 cells (examining the effect of oxidative stress alone), unstressed and stressed in WT U-2 OS cells (examining the effect of stress together with SGs) and stressed ΔΔG3BP1/2 cells and stressed WT U-2 OS cells (examining the effect of SGs alone), we identified only a few nucleotide bases within the *AHNAK* 3′UTR that exhibited statistically significant changes in DMS reactivity (Fig. 5H-J).

Thus, although RNA accessibility to chemical probing may change during oxidative stress and upon SG formation, these changes are not widespread but instead are limited to a small number of isolated bases distributed throughout the *AHNAK* 3′UTR.

### DMS-MaPseq probing revealed modest changes in base accessibility of domains 3 and 4 of *NORAD* lncRNA during oxidative stress

To determine whether observations made with *AHNAK* 3′UTR extended to other SG-RNAs, we also probed the base accessibility of lncRNA *NORAD* to DMS. Specifically, we focused on the 1332 nt region previously annotated as *NORAD* domains 3 and 4 (termed ND3 and 4; Fig. 6A) (Matheny et al., 2021). *NORAD* domains 1 to 5 are repetitive and highly conserved (Lee et al., 2016). Therefore, we reasoned that probing ND3 and ND4, which together represent approximately 50% of the *NORAD* transcript, could provide insight into the overall accessibility of the repetitive *NORAD* domains 1–5.

**Figure 6:**
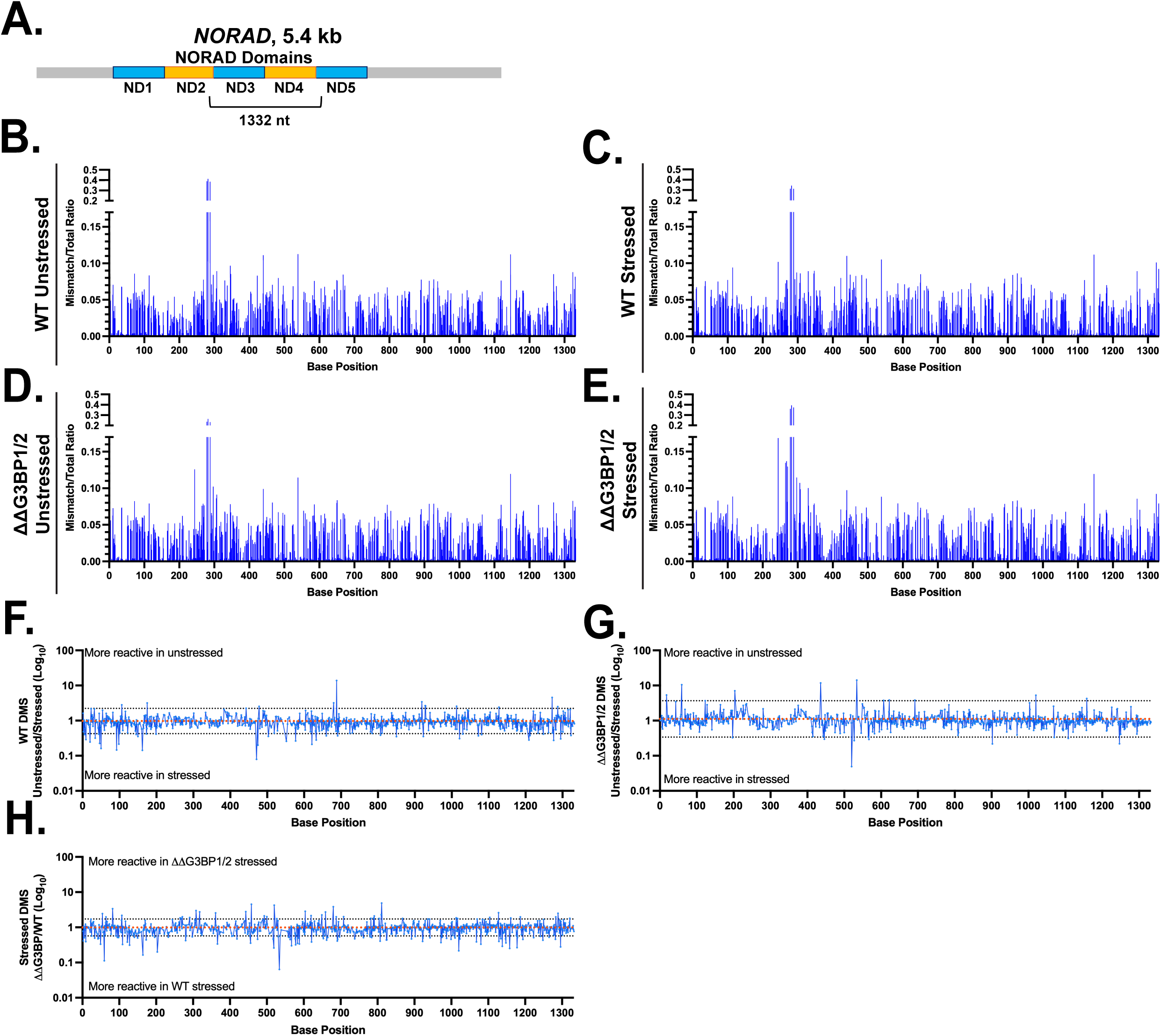
DMS-MaPseq probing revealed limited changes in base accessibility of domains 3 and 4 of *NORAD* lncRNA during oxidative stress. A. Schematic of the lncRNA *NORAD* with the position of its 1332 nt region assayed with DMS-MaPseq. B-E. DMS mismatch ratio of the *NORAD* ND3 and ND4 region in unstressed (B) and stressed WT cells (C) in in unstressed (D) and stressed ΔΔG3BP1/2 (E) cells. The mismatch ratio represents the ratio of reads with DMS modification-induced mismatches compared to the reference transcriptome. The ratio shown is an average of three biological replicates. F-H. DMS ratios in the average per base DMS reactivity of the *NORAD* ND3 and ND4 region in unstressed and stressed WT U-2 OS cells (F), unstressed and stressed ΔΔG3BP1/2 U-2 OS cells (G) and stressed ΔΔG3BP1/2 and stressed WT U-2 OS cells (H). The orange dotted lines denote the average ratio between two compared conditions while the black dotted lines represent the statistical significance threshold where bases that fall above and below the line show significant differences between the conditions examined. The ratio is calculated from the averages of the three biological replicates per condition (see Materials and Methods).

As with *AHNAK* 3′UTR, we observed a high correlation of the average base accessibility to DMS for ND3 and ND4 across biological replicates, ranging from 0.87 to 0.97 in unstressed and from 0.92 to 0.96 in stressed cells (Fig. S10A,B). Similarly, the majority of bases within *NORAD* ND 3 and ND4 did not display global changes in DMS reactivity in response to G3BP1/G3BP2 expression or arsenite treatment (Fig. 6B-E). Instead, only a small number of bases displayed statistically significant changes in DMS reactivity across these conditions (Fig. 6F-H).

Therefore, similar to the *AHNAK* 3′UTR, our data suggest that neither oxidative stress nor expression of G3BP1/2 substantially alters the accessibility of *NORAD* ND3 and ND4 to chemical probing.

## Discussion

In this study, we examined intermolecular base pairing during oxidative stress in U-2 OS cells using AMT crosslinking and reporter RNAs. As proof of principle, we showed that AMT crosslinking captures known base pairing interactions between the U3 snoRNA and the pre-18S rRNA, which is required for processing of the mature 18S rRNA. This result establishes two things about the assay. First, it captures base-pairing interactions even within a protein-RNA condensate, since the U3 snoRNA–pre-18S rRNA interaction occurs within the nucleolus. Second, this interaction persisted during arsenite-induced oxidative stress, indicating that it is robust to this stress condition. Different cellular environments will likely affect intermolecular RNA:RNA interactions in distinct ways, but this result establishes that protein-RNA-rich environments do not, on their own, block AMT’s ability to detect intermolecular base pairing interactions.

Ratiometric enrichment, as used here to study intermolecular base pairing, is not an absolute measure of interaction frequency and may be influenced by the efficiency of reversal of AMT-induced crosslinks, RNA fragmentation during UV treatment and purification, and the efficiency of PCR amplification after precipitation (see also Materials and Methods) (Aw et al., 2016, Zhang et al., 2021) It therefore reports on the levels of RNAs that could be crosslinked with AMT, by virtue of being accessible to AMT and having nucleotides in sufficient proximity to crosslink, and does not reflect absolute levels. For instance, although our detection of the U3 snoRNA–pre-18S rRNA interaction is robust within the nucleolus, the efficiency of detection may have been affected by the protein and RNA density of the condensate and the stress condition, preventing us from examining changes in the abundance of this interaction in response to cellular perturbations in finer detail. With this caveat, the 0.35 ratiometric enrichment we measured between U3 snoRNA and pre-18S rRNA (Fig. 1D) offers a useful reference point for the ratiometric enrichment expected for RNA pairs whose function depends on intermolecular base pairing. Ratiometric enrichment of our 3SL sense:antisense reporter mRNAs, fused with MS2 repeats fall below this reference point: 0.16 during arsenite stress in cells expressing endogenous G3BP1/2 (Fig. 3D), falling further to 0.02 upon co-expression of MCP-G3BP1 (Fig. 3F). Therefore, in this case, the ratiometric enrichment suggests that intermolecular base pairing between the MS2-tagged 3SL sense:antisense luciferase reporters during oxidative stress is not as high as that observed for the U3 snoRNA–pre-18S rRNA pair, and is modulated by G3BP1 expression.

Changes in ratiometric enrichment may suggest regulation that may involve RNA helicases or other RBP regulators, but they can also result from changes in RNA levels (locally or cell-wide), the absence of proteins that could augment intermolecular base pairing, or the lack of RNA sequences that promote such interactions. Indeed, we observed increased detection of interactions between MS2-tagged luciferase reporters that depended on both co-expression of MCP-G3BP1 and the presence of RNA sequences that promote base pairing; these sequences alone generated sufficient co-precipitation of in vitro transcribed luciferase reporters even in the absence of protein or AMT (Fig. 2C).

This RNA sequence- and protein-driven intermolecular base pairing has been observed in other systems. The *Drosophila oskar* mRNA dimerizes and oligomerizes in vivo through the combined action of RNA concentration, RBPs, and an RNA palindrome capable of base pairing (Ye et al., 2026, Jambor et al., 2011). Similarly, RNA viruses such as Influenza A, Human immunodeficiency virus-1 (HIV-1), Murine leukemia virus (MLV), and Flock House virus (FHV) rely on intermolecular base pairing to organize and package their RNA genomes (Berkowitz et al., 1996, Dadonaite et al., 2019, De Tapia et al., 1998, Johnson and Telesnitsky, 2010, Kim and Tinoco, 2000, Krishna and Schneemann, 1999, Le Sage et al., 2020, Paillart et al., 2004, Zhou and Routh, 2024), a process influenced by high RNA concentration and co-expression of viral proteins (Maurel and Mougel, 2010, Muriaux et al., 1996, Weiss et al., 1984). Even our proof-of-principle interaction fits this pattern: U3 snoRNA and pre-18S rRNA base pair through evolutionarily conserved complementary sequences, assembled by conserved RBPs into the protein-RNA complex that carries out pre-18S rRNA processing (Beltrame and Tollervey, 1995).

### Modest spatial overlap between *AHNAK* mRNA and lnc *NORAD*

The limited sensitivity of AMT crosslinking prevented us from examining interactions between *AHNAK* and *NORAD* in cells. However, smFISH coupled with co-localization measurements revealed that *AHNAK* mRNA and lncRNA *NORAD* exhibited modest spatial overlap both in unstressed and stressed WT cells (Fig. 4C), an observation that applied to other SG-enriched RNA pairs (Fig. S8K,L). smFISH cannot resolve intermolecular base pairing directly, but the modest increase in overlap during stress suggests that interactions between these RNAs, whether base pairing or protein-driven, were not abundant enough to cause extensive spatial mixing of the two transcripts. This, along with the limited AMT sensitivity, could also contribute to our inability to detect base pairing between these RNAs with AMT.

A scarcity of intermolecular interactions would not make them unimportant. If each transcript forms only a few base-pairing contacts at a time, but does so with many different partners across the population, the resulting multivalency could still organize substantial numbers of RNAs. Such multivalency could support SG assembly while also enabling their rapid disassembly once stress resolves, since fewer individual contacts would need to be broken. Consistent with a functional role for G3BP1 here, loss of endogenous G3BP1/2 leaves RNAs dispersed during stress (Fig. 4H,I; S8G-J). In agreement with this result, the presence of endogenous G3BP1/2 enhances AMT-dependent detection of intermolecular base pairing of the 3SL sense:antisense reporter pair during stress (Fig. 3D). Given that G3BP1 engages in multivalent protein-protein and protein-RNA interactions and, in vitro, facilitates intermolecular RNA:RNA interactions (Parker et al., 2025, Trussina et al., 2025, Yang et al., 2020), our result further supports the idea that in cells, stress-induced intermolecular RNA:RNA interactions are mediated by the presence of endogenous G3BP1/2.

### G3BP1 may have a dual, stress-dependent role in modulating intermolecular base pairing

Our measurements cannot establish whether interactions among 3SL sense:antisense reporter mRNAs occur in the cytosol or within SGs during oxidative stress. However, they reveal an interesting reversal in the effect of G3BP1. Ectopic MCP-G3BP1 expression increased ratiometric enrichment of the luciferase reporters in unstressed cells, yet suppressed this same enrichment once arsenite stress was applied (Fig. 3F). Meanwhile, WT U-2 OS cells lacking ectopic MCP-G3BP1 showed the opposite trajectory, with enrichment rising only during stress (Fig. 3D).

This reversal argues against a simple model of G3BP1 as a promoter of base pairing. Instead, our results suggest that G3BP1 may act as a potential switch whose effect depends on cellular stress state: at endogenous levels, G3BP1 may favor base pairing during stress, while at elevated levels, it instead may drive precocious pairing before stress and curtail it afterward. Although the underlying mechanism remains undefined, this behavior parallels in vitro and in vivo work showing that G3BP1 both promotes RNA:RNA interactions, by concentrating RNAs into proximity, and restrains them, potentially via recruitment of DEAD-box RNA helicases (Parker et al., 2025, Trussina et al., 2025, Tauber et al., 2020). Therefore, our data, together with these published observations, suggest a possible dual role for G3BP1 in the formation of intermolecular base pairing interactions, acting as both a promoter and an attenuator. In support of this dual role, a recent report found that G3BP1 both enhances and suppresses translation in a manner that reinforces the stress condition program (Smith and Bartel, 2026).

### DMS-MaPseq shows modest changes in base accessibility of specific SG-enriched RNA regions

Our DMS-MaPseq results on base accessibility within the 3′UTR of *AHNAK* and the ND3/ND4 domains of *NORAD* show similar DMS reactivity profiles across stress treatment and in the absence of G3BP1/2, with only a few bases displaying significant fold changes between these conditions (Fig. 5, 6). A recent study found that, during oxidative stress, 3′UTRs across the transcriptome undergo less structural remodeling than 5′UTR and CDS regions, and that SG-enriched RNAs undergo less remodeling than non-SG-enriched mRNAs (Farenhem et al., 2025, Zhu et al., 2025, Wu et al., 2024), a pattern attributable to translational inhibition rather than to SG localization per se (Farenhem et al., 2025). The tendency of RNAs to minimize changes in single-strandedness within SGs (Zhu et al., 2025) may reflect the fact that SG-enriched RNAs are already highly structured prior to stress (Farenhem et al., 2025). This is consistent with the observation that RNAs enriching well in SGs are less translationally efficient in unstressed cells than those enriching poorly (Khong et al., 2017), supporting a model in which SG-enriched RNAs are less subject to translation-repression-mediated structural remodeling. This observation is also consistent with our previous work on *Drosophila*, where intermolecular base pairing among 3′UTR of germ granule mRNAs is constrained by RNA structure (Tian et al., 2025). We do not know whether the overall maintenance of DMS base accessibility reflects intramolecular base pairing, intermolecular base pairing among mRNAs, or protein binding. However, our measurements reported here, together with published observations (Tian et al., 2025, Farenhem et al., 2025, Wu et al., 2024, Zhu et al., 2025), suggest that minimal structural rearrangement may more broadly reflect the regulation of intra- and intermolecular base pairing within protein-RNA condensates.

Collectively, our data show that the abundance of intermolecular base pairing in cells undergoing oxidative stress is influenced by endogenous G3BP1/2, exogenous MCP-G3BP1, and RNA sequence (Fig. 7). While these interactions may contribute to the multivalency required for SG stabilization, their limited abundance would allow SGs to remain dynamic, enabling rapid dissolution once stress passes, a process further facilitated by RNA regulators such as RNA helicases (Tauber et al., 2020, Trussina et al., 2025).

**Figure 7:**
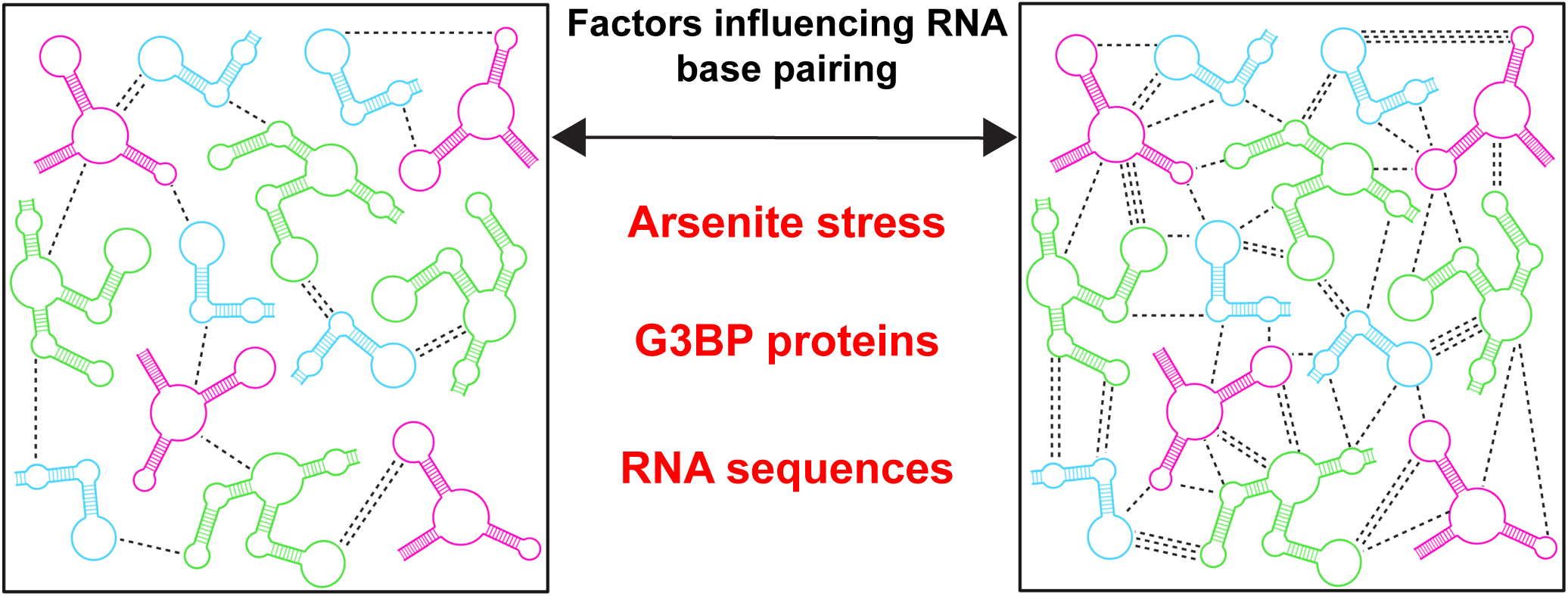
Intermolecular base pairing among RNAs is modulated in mammalian cells during oxidative stress. Schematic showing how RNA sequence, G3BP1/2 proteins, and oxidative stress modulate the detection of intermolecular base pairing. This modulation may act together with RNA structure to control the abundance of intermolecular base pairing and preserve RNA function during oxidative stress.

### Limitations

Our study shows that AMT crosslinking can be applied to examine intermolecular base pairing interactions in cellulo. While this approach is successful, several limitations curtail its sensitivity. Apart from the technical limitations discussed above (crosslink reversal, fragmentation, and PCR bias), AMT crosslinking efficiency itself could vary systematically between RNA species (se also Materials and Methods). If base pairing-prone sequences within a given RNA are few, AMT-dependent co-precipitation would be correspondingly low, underestimating the true abundance of intermolecular interactions for that RNA. Because RNA structure can occlude interaction-prone sequences, as our work in *Drosophila* revealed (Tian et al., 2025), it is feasible that different cellular environments, including condensation promoted by G3BP1/2, alter RNA structure enough to expose or occlude such sequences. Our data are consistent with the idea that AMT detects interactions more robustly for RNAs that contain interaction-prone sequences or are bound by proteins that augment such interactions, and possibly for more highly expressed RNAs. Detection for transcripts lacking these features would be noisier, making it harder to examine changes in the abundance of their interactions across cellular conditions. Together, these factors limit the sensitivity and interpretability of AMT crosslinking for detecting intermolecular base pairing in cells.

## ACKNOWLEDGEMENTS

We would like to thank the members of the Trcek lab and Drs. Philip C. Bevilacqua, Anthony Leung, John Kim, and Mollie Meffert for the critical reading of our manuscript. We would also like to thank Siran Tian and Aritra Basu for their initial help with in vitro RNA transcription and image segmentation and Dr. J. Paul Taylor for sharing their ΔΔG3BP1/2 U-2 OS and G3BP1:GFP-expressing cells with us and Dr. Leung for sharing the CI-neo-His-MS2BP-G3BP1-WT plasmid with us.

## COMPETING INTERESTS

The authors declare no competing interests.

## FUNDING

This research was supported by the NIGMS R35GM142737 grant awarded to TT.

## DATA AND RESOURCE AVAILABILITY

Microscopy data generated in this study will be shared by the lead contact upon request. This paper does not report original code. Any additional information required to reanalyze the data reported in this paper is available from lead contact upon request. Further information and requests for resources and reagents should be directed to and will be fulfilled by the lead contact, Tatjana Trcek.

## AUTHOR CONTRIBUTION

TT wrote the initial draft of the manuscript with JL providing the methods section, legends and manuscript edits. JL generated all the figures. JL and TT: conceptualization. JL designed and performed the majority experiments with IB and ES, helping with the investigation and analysis.

## MATERIALS AND METHODS

### Mammalian cell culture

Human U-2 OS cells were obtained from the American Type Culture Collection (ATCC; HTB-96). The U-2 OS G3BP1:GFP and U-2 OS ΔΔG3BP1/2 cells lines were graciously gifted to us by Paul Taylor’s lab (Yang et al., 2020, Zhang et al., 2018). Cells were cultured in DMEM (Corning; 10-013-CV) supplemented with 10% Fetal Bovine Serum (FBS; ThermoFisher Scientific; A4736201), 100U/mL penicillin and 100 μg/mL streptomycin (Millipore Sigma; P4333), within a 37°C humidified incubator with 5% CO_2_. Stable U-2 OS cell lines expressing TET3G and 6x-His-MCP-G3BP1 were cultured in background concentrations of supplemented 200ug/mL G418 sulfate (Millipore Sigma; G8168) and 100ug/mL Hygromycin (Invivogen; ant-hg-1) in complete DMEM, respectively. All cell line experiments were completed with cultured cells under 30 passages.

### Transfection of Cells

Stable transfection of TET3G or 6xHis-MCP-G3BP1 plasmid constructs into cell was performed using Xfect Transfection Reagent (Takara; 631318). Transfections were performed according to manufacturer protocols. Briefly, cells were seeded into 6-well plates and transfected with 2ug of either pEF1α-TET3G plasmid containing a G418 resistance gene or the pCI-neo-intron-His-MS2BP-G3BP1-WT plasmid co-transfected with a Hygromycin linear marker DNA (Takara, 631625), which conferred resistance to Hygromycin. After 4 hours (hrs) of transfection, cells were split into four 10 centimeters (cm) dishes and left to recover for 48 hrs. Cell clones were selected with complete DMEM supplemented with 500ug/mL of G418 or 200ug/mL Hygromycin, which was replaced every 4 days for ∼2 weeks. Resistant clones were isolated and screened by qPCR, immunofluorescence, and/or western blot. For transient transfections of the RLuc/FLuc reporters, cells were transfected with 7.5ug of the pTRE3G-FLuc-sense-12xMS2_pTRE3G-RLuc-Antisense-12xMS2 plasmid in a 6-well plate, scaled up to 45ug of plasmid for 10 cm plate transfections. Cells were then treated with 1ug/mL of doxycycline hyclate (Millipore Sigma; D9891) in DMEM at 37⁰C for 36 hours. The doxycycline-containing media was replaced every 24 hrs.

### Induction of stress condition

U-2 OS cells undergoing stress treatment were cultured as described above. To induce oxidative stress, the cells were treated with 500μM sodium arsenite (NaAsO_4_; Millipore Sigma; S7400) for 1 hour at 37⁰C.

### DNA Constructs

DNA plasmids were generated using Gibson assembly with NEB HiFi DNA Assembly mastermix (NEB; E2621) or restriction digest followed by T4 DNA ligation (NEB; M0202). All DNA plasmids were confirmed by whole plasmid nanopore sequencing.

### Design of the Firefly/Renilla reporter mRNAs

We employed the *FLuc* and *RLuc* luciferase reporter mRNAs fused with the 3′UTR of the *Drosophila shutdown* (*shu*) gene. Into each reporter, we inserted three stems derived from the SL1 motif required for dimerization of HIV RNA (Dirac et al., 2002, Paillart et al., 1996, St Louis et al., 1998), as we have done previously (Tian et al., 2025). AMT preferably crosslinks pyrimidine dinucleotides on adjacent RNA strands (Calvet and Pederson, 1979, Cimino et al., 1985, Nilsen, 2014, Stadler and Stafforst, 2014, Velema and Lu, 2023). To account for this preference, the 3′UTR of the 3SL sense:antisense reporter contained AUUAUA sequences in two SL1 loops and a CGGCGG sequence in the third SL1 loop within the *Firefly* 3′UTR, and exposed UAUAAU sequences in two SL1 loops and a CCGCCG sequence in the third SL1 loop within the *Renilla* 3′UTR (Fig. 2A) as we have done previously (Tian et al., 2025). For in cellulo experiments, these 3′UTRs were each fused with twelve MS2 bacteriophage-derived hairpins (Bertrand et al., 1998) (Fig. 3A, Fig. S5A). To ensure balanced expression of *FLuc* and *RLuc* mRNAs within the same cell during transient transfection, we utilized a plasmid in which both reporters were driven by two sets of the same tetracycline-inducible promoter (pTRE3G Vector Set; Takara; 631173).

### Western Blotting

Cells were detached with 1ml of Trypsin-EDTA (0.25%; Thermo Fisher Scientific, 25200056) for 5 min, quenched with DMEM, then washed with ice-cold 1x PBS. Cells were then lysed with RIPA buffer (25 mM Tris•HCl pH 7.6, 150 mM NaCl, 1% IGEPAL CA-640, 1% sodium deoxycholate, 0.1% SDS). Afterwards, the protein supernatant was isolated by centrifugation and quantified by a BCA assay (ThermoFisher Scientific; 23225). 20µg of protein was diluted with 4X Laemmli Sample Buffer (Bio-Rad; 1610747) to a final concentration of 1X, then boiled at 95⁰C for 5 min. Samples were then loaded onto 7.5% Mini-PROTEAN TGX gels (Bio Rad, 4561023) with Precision Plus Dual Color Standards (Bio Rad, 1610374). Proteins were then transferred onto PVDF membranes using semi-dry conditions with assembled transfer sandwiches equilibrated in 1X Towbin transfer buffer (25 mM Tris, 192 mM glycine, 20% (v/v) methanol (pH 8.3)) and the Transblot TURBO system (Trans-Blot® Turbo™ Transfer System, Bio Rad, 1704150). PVDF membranes were blocked with 5% milk in 1XTBST (20mM Tris, 150mM NaCl, 0.1% w/v Tween-20), then incubated with primary anti-G3BP1 antibody (Abcam, ab181149) at 1:10000 dilution or anti-GAPDH (Abcam, ab181602) at 1:10000 in the 5% milk blocking buffer at 4⁰C overnight (O/N). To visualize protein bands, membranes were then incubated with HRP-conjugated Goat anti-rabbit IgG secondary antibodies (Abcam; ab6721) at 1:10000-1:20000. Bands were detected with chemiluminescence using SuperSignal West Dura Extended Duration Substrate (ThermoFisher Scientific; 34075) and imaged with the Bio-Rad ChemiDoc MP Imager.

### RNA isolation and RT-qPCR

RNA from U-2 OS cells was extracted with Trizol (ThermoFisher Scientific; 15596026) and chloroform (Fisher Scientific; BP1145-1) following manufacturer protocols. 3μg of total RNA was treated with RQ1 RNase-free DNase (Promega: M6101). cDNA synthesis was performed 1 μg of RNA using SuperScript III reverse transcriptase (Thermo Fisher; 18080093) using gene-specific primers or random hexamers following manufacturer protocols (Table S2). qPCR was performed using ssoSYBR Advanced SYBR Green Mastermix (Bio-Rad; 1725272) in a 20uL reaction using 1 μL each of 10 μM forward and reverse primers (Table S2). qPCR samples were run either with *GAPDH* for relative quantification, or with standard curves for absolute quantification. For absolute quantification, qPCR samples for specific RNA targets were run alongside standard curves generated from 10-fold dilutions of the same qPCR amplicons that were separately amplified by Q5 HiFi DNA polymerase (New England Biolabs; M0492S) and extracted from agarose gels using Monarch Gel Extraction kit (New England Biolabs; T1120L). These standard curves were used to determine the number of transcripts detected per ng of RNA input. Quantitative PCR analysis was performed using a CFX Opus 96 Real-Time PCR System from Bio-Rad. The primer sequences are listed in Table S2.

### In-house labeling of smFISH probes

smFISH probe sequences were designed using Stellaris Probe Designer with a minimum of 24 probes per RNA target (Table S3). The oligos were ordered from IDT and fluorescently labeled in-house using terminal deoxynucleotidyl transferase (Thermo Fisher; EP0161) and amino-11-ddUTP (Lumiprobe; 15040), followed by covalent conjugation with AF568 (Lumiprobe: 24820) or AF647 (Lumiprobe: 26820) NHS esters as previously described (Gaspar et al., 2017, Tian et al., 2025). The sequences of smFISH probes are listed in Table S3.

### smFISH in U-2 OS cells

U-2 OS cells were fixed and hybridized with smFISH probes as described before (Trcek et al., 2013).

### Immunostaining of U-2 OS Cells

U-2 OS cells, grown on collagen coverslips, were fixed and permeabilized as described above. To block non-specific antibody staining, the cells were then incubated with 1% Bovine Serum Albumin (BSA; Millipore Sigma; A6003) /0.1%Triton X-100/1x PBS blocking buffer for 15 min. Cells were then incubated with primary antibodies at 4⁰C O/N. The following primary antibodies were used: G3BP1 (1:500, rabbit, ab181149), TIA1 (1:200, mouse, sc-48371), CAPRIN1 (1:500, rabbit, 15112-1-AP), USP10 (1:200, rabbit, 19374-1-AP), ATXN2 (1:250, mouse, 611387), 6xHis (1:200, mouse, MA1-135). After primary antibody incubation, and subsequent 1X PBS washes (3 washes, 10min each at RT), cells were incubated with secondary antibodies in blocking buffer. The following secondary antibodies were used: Goat Anti-Rabbit IgG CF640R (1:1000, SAB4600399), Goat Anti-Mouse IgG AF647 (1:1000, A-21235), Goat Anti Rabbit IgG AF488 (1:1000, ab150077). After primary antibody incubation, and subsequent 1X PBS washes, cells were incubated with secondary antibodies in blocking buffer. The following secondary antibodies were used: Goat Anti-Rabbit IgG CF640R (1:1000, SAB4600399), Goat Anti-Mouse IgG AF647 (1:1000, A-21235), Goat Anti Rabbit IgG AF488 (1:1000, ab150077). Afterwards, cells were washed 3-times 1X PBS for 5 min each at RT. Nuclear DNA was stained with DAPI and the coverslips were mounted using Prolong Glass as described above.

### Immunofluorescence coupled with smFISH

U-2 OS cells were grown on collagen-coated coverslips (Neuvitro, GG-12-Collagen), washed twice with 1xPBS, fixed with 4% paraformaldehyde (PFA; Electron Microscopy Sciences; 15713), then washed three times with 1x PBS. The cells were then permeabilized with permeabilization/blocking (PB) buffer containing 1X PBS, 1% acetylated-albumin (Millipore Sigma; B2518), 0.3% Triton X-100 (Millipore Sigma; T8787), 2mM Ribonucleoside vanadyl complex (VRC; New England Biolabs; S1402S) for 1 hr at room temperature (RT), then incubated with G3BP1(1:500, ab181149) primary antibodies diluted according to manufacturer recommendations in PB buffer at 4⁰C O/N. Cells were then washed 3 times with 1X PBS then incubated with F(ab’)2-Goat Anti-Rabbit IgG (AF488) (1:1000, A11070) secondary antibodies diluted according to manufacturer recommendations in PB buffer for 90 min at RT. The samples were washed with 1X PBS, then fixed again with 4% PFA. The cells were then hybridized with smFISH probes as described before (Trcek et al., 2013). After smFISH hybridization, the nuclear DNA were counter-stained with 1ug/mL 4’,6-Diamidine-2’-phenylindole dihydrochloride (DAPI; Millipore Sigma; 10236276001) and mounted on glass slides (Fisher Scientific, 155442) using ProLong Gold Antifade Mountant (ThermoFisher Scientific; P36930).

### Microscopy and deconvolution

Images were acquired in three dimensions (3D) with a vt-instant Structured Illumination Microscope (vt-iSIM; BioVision Technologies) equipped with the 405nm 100mW, 488nm 150mW, 561nm 150mW, 642nm 100mW, and 445nm 75 mW lasers, two ORCA-Fusion sCMOS cameras and the Leica HC PL APO 63x/1.30 GLYC CORR CS2, HC PL APO 63x/1.40 OIL CS2 and HC PL APO 100x/1.47 OIL CORR TIRF objectives as described before (Curnutte et al., 2023). Images were then deconvolved using Huygens (Scientific Volume Imaging) as we have done before (Tian et al., 2025).

### 4′ AMT Crosslinking and Extraction of in cellulo RNA

Confluent (∼80%) 10 cm cell culture plates of U-2 OS cells washed with twice 1xPBS, then treated with 1mL of 1μg/mL AMT (Millipore Sigma; A4330) for 15 min at 37°C. The plates were transferred onto ice and irradiated in a UV crosslinker (UVP CL-1000L Longwave UV Crosslinker, 95-0228-01) using 365 UV irradiation to induce RNA crosslinking (Fig. 1B) at two-10 min intervals for a total of 20 min with plate lids off, swirling between 10 min intervals. The AMT was then removed and cells were washed twice with 1x PBS. Cells were processed using the total nucleic acid extraction (TNA) protocol to extract RNA, as described before (Zhang et al., 2021). In short, cells were lysed with 100uL of 6M Guanidine thiocyanate (Millipore Sigma; 50983), scraped, and collected in microfuge tubes. 12μL of 500mM Ethylenediaminetetraacetic acid (EDTA; Fisher Scientific; 15-575-020) and 60uL of 10xPBS were added to the samples and the volume was brought up to 600uL using nuclease-free water (Thermo Fisher Scientific, AM9937). Insoluble material was broken up by passing the sample through the 25G needle. Proteinase K (New England Biolabs; P8107S) digestion was performed at 37⁰C for 1hr on a heated shaker. After digestion, 60uL of 3M sodium acetate (Fisher Scientific; AM9740) and 600uL of water-saturated phenol (ThermoFIsher Scientific; 15513039) were added to the sample, then split evenly between two tubes. To precipitate the nucleic acids, 600uL of isopropanol (Fisher Scientific; A416-1) was added and the sample and the tubes were spun at 15000 rotations per minute (rpm) for 20 min at 4°C. The nucleic acid pellet was washed three times with 70% ethanol and resuspended in 150uL of nuclease-free water. RNA concentration within samples was measured by Nanodrop. Finally, to purify RNA from the nucleic acid sample, 10-100ug of TNA samples were digested with 50 units of Turbo DNase (ThermoFisher Scientific; AM2238). Samples were the precipitated using the TNA method described above and eluted in 50uL of nuclease-free water. Purified RNA was then measured again with Nanodrop.

### AMT RNA enrichment and qPCR of *in cellulo* samples

After RNA extraction and removal of DNA and interacting proteins, the AMT-crosslinked RNA was precipitated using biotinylated DNA oligonucleotides following an established protocol (Aw et al., 2016). Specifically, 3-10μg of purified, AMT-crosslinked RNA was mixed with 1uL of 100uM (100pmol total) of biotinylated probes (IDT) and diluted to a final volume of 300uL with water. 600uL of fresh hybridization buffer (750mM NaCl (ThermoFisher Scientific; AM9759), 1% SDS, 50mM Tris (ThermoFisher Scientific; AM9855G), 1mM EDTA, 15% formamide (Millipore Sigma; 344206-M)) supplemented with SUPERase-In RNase Inhibitor (ThermoFIsher Scientific; AM2694) (1:200) was then added to the samples and left to hybridize at 37°C O/N with shaking (Eppendorf® New Brunswick™ Innova® 40, 80rpm). The tubes were taped in an empty pipette tip box sprayed with RNAse Zap. 50uL of Streptavidin beads (Dynabeads MyOne Streptavidin C1; ThermoFisher Scientific; 65001) were prepared for each RNA sample following the manufacturer guidelines, then mixed with the hybridization sample at RT for 15 min with inversion (Variable Speed Tube Revolver with Digital Display TR-02UA, 10rpm). The beads were then washed 5 times with 37°C wash buffer (0.1X SCC (ThermoFisher Scientific; AM9763), 0.5% SDS), eluted using 100μg of Proteinase K with 95uL of PK Buffer (100mM NaCl, 10mM TrisCl PH 7.0, 1mM EDTA, 0.5% SDS) and incubated at 50°C for 45 min in Thermoshaker at 500rpm, followed by boiling at 95⁰C for 10 min. Samples were purified using the TNA method described above and resuspended in nuclease-free water in 50uL. The eluted RNA was reverse crosslinked with a 254nM crosslinker (AnalytikJena CX-2000 UV crosslinker, AJ 95-0339-01) and further purified using the Zymo RNA Clean and Concentrator-5 Kit (Zymo Research; R1014). The RNA samples were then reversed transcribed with SuperScript IV Reverse Transcriptase (ThermoFisher Scientific; 18090010) using gene specific primers (Table S2). qPCR was performed using SsoSYBR Advanced SYBR Green Mastermix and CFX Opus 96 Real-Time PCR System.

### Co-precipitation of the AMT-crosslinked and *in vitro* transcribed RNA

Following our previous work (Tian et al., 2025), RNAs were *in vitro* transcribed by MEGAscript T7 transcription kit (ThermoFisher Scientific; AM1334) and recovered by Trizol (ThermoFisher Scientific; 15596026) extraction and isopropanol (Fisher Scientific: A416-1) precipitation. The resulting RNA pellets were washed with 80% ethanol and resuspended in nuclease-free water (ThermoFisher Scientific: AM9937). The In vitro RNA pulldowns were done using a published protocol (Bose et al., 2024) with minor modifications. 20pmol of *in vitro* transcribed *RLuc* transcripts (CDS or CDS + 3′UTR) and 10pmol of *FLuc* transcripts (CDS or CDS + 3′UTR) were used. *RLuc* and *FLuc* transcripts were individually heated up to 95⁰C for 2min, followed by incubation on ice for 15min. The transcripts were incubated separately in a high salt dimerization buffer to a final concentration of 50mM sodium cacodylate, 300mM KCl and 5mM MgCl_2_, at 37⁰C for 30min, before mixing together and incubating at 37⁰C for an additional 30min. Then AMT (or water) was added to the mixed samples for a final concentration of 50ng/uL and incubated at 37⁰C for 15 min before 365nm UV crosslinking. Finally, the same co-precipitation protocol was used as described previously, but with half the hybridization reaction volume and half the amount of streptavidin beads used for pulldown.

### Dimethyl-Sulfate (DMS) Treatment of RNAs

Targeted DMS-MaPseq of the *AHNAK* 3′UTR and *NORAD* domains 3 and 4 were performed previously described (Tomezsko et al., 2020, Zubradt et al., 2017). Briefly, cells were treated with 1% Dimethyl sulfoxide (DMS; Millipore Sigma; 276855) in DMEM at 37⁰C for 5 min with gentle swirling (Eppendorf New Brunswick Innova Incubator Shaker, 50rpm). The reaction was then quenched with 30% v/v 2-mercaptoethanol (Millipore Sigma, M3148), after which the RNA was extracted with Trizol and cleaned with Zymo RNA Clean and Concentrator Kit. The extracted RNA was DNase treated (Thermo Fisher, AM2238) and rRNA depleted using rRNA subtraction oligos (Table S5), as previously described (Adiconis et al., 2013) and RNase H (New England Biolabs; M0297S) following manufacturer specifications. 50ng of RNA was reverse transcribed with Induro reverse transcriptase (New England Biolabs; M0681S) using pooled gene specific primers at 5pmol each (Table S4). The reverse transcription was performed at 55⁰C for 2 hr. The RNA regions selected for probing with DMS were split into overlapping fragments of approximately 150-200bp in length and PCR amplified using Q5 high-fidelity polymerase. The amplified fragments were run on 2% agarose gels with 1X SYBR safe (Thermo Fisher; S33102) and extracted using Monarch Spin DNA Gel Extraction Kit (New England Biolabs; T1120L).

### DNA library preparation and sequencing

The concentration of each DNA fragment was determined using Qubit 4 fluorometer with the 1X dsDNA HS assay kit (Thermo Fisher; Q32854). 10ng of each fragment was pooled together into a mixture that contained a total of 140ng of DNA fragments per RNA. Sequencing libraries were prepared using NEBNext Ultra II DNA library prep kit (New England Biolabs; E7645S) for Illumina sequencing. Libraries were sequenced on the MiSeq system by the Johns Hopkins Genetic Core Research Facility.

### Analysis of RNA base pairing and ratiometric enrichment of AMT treated samples

RNA samples were separately hybridized with a biotin probe set specific to an RNA of interest as well as a biotin probe set against the GFP sequence as was previously published (Aw et al., 2016). Since the GFP sequence is not expressed, GFP probe hybridization serves as a background pulldown control, allowing quantification of fold enrichment for both the target RNA and any interacting RNA sequences (Fig. S1A). Validation of this approach against known RNA base-pairing partners, the pre-18S rRNA and U3 snoRNA, showed detectable fold enrichment of U3 snoRNA relative to background even in the absence of AMT crosslinking (Fig. S1B,C). Treatment of cells with AMT increased Cq values and correspondingly decreased fold enrichment of the target pre-18S rRNA (Fig. S1B,C), which may have resulted from AMT crosslinking-induced RNA damage, inefficient reversal of crosslinks, and reduced biotin-probe accessibility (Aw et al., 2016, Zhang et al., 2021). Nevertheless, AMT treatment increased U3 snoRNA fold enrichment despite the decrease in pre-18S rRNA fold enrichment, suggesting an increase in detectable base pairing interactions through crosslinking (Fig. S1C). To account for these factors, we calculated ratiometric enrichment by dividing the fold enrichment of a co-precipitated RNA by the fold enrichment of the target RNA hybridized with its specific biotin probe set (Fig. S1A). Ratiometric enrichment therefore reports the proportion of an interacting RNA co-precipitated per target RNA, while accounting for background pulldown levels of both target and interacting RNAs, as well as for the factors by which AMT crosslinking influences the amount of RNA detected. This works well for highly abundant interacting RNA pairs (Fig. 1D).

However, the sensitivity of ratiometric enrichment decreases when pulldown detection and fold enrichment of the interacting RNA become very low. In this case, ratiometric enrichment can become driven by the pulldown efficiency of the target RNA rather than by base pairing between the two RNAs, as was the case for *AHNAK* and *NORAD* (Fig. S7F,G). For example, in ΔΔG3BP1/2 cells, *NORAD* amplicon 4 shows significant ratiometric enrichment during arsenite stress relative to cells not treated with AMT (Fig. S7H). However, fold enrichment of this amplicon is close to zero across all conditions (Fig. S7I). The significant increase in ratiometric enrichment is therefore driven by a decrease in fold enrichment of the target *AHNAK* mRNA across conditions, rather than by base pairing between *AHNAK* and *NORAD* in this region. This demonstrates that AMT crosslinking has limited sensitivity for detecting intermolecular base pairing when the underlying interactions are of low abundance.

### Analysis of DMS-MaPseq data

The raw fasta files were analyzed using the Detection of RNA folding Ensembles using Expectation-Maximization (DREEM) as previously described (Tomezsko et al., 2020) and implemented in the RNA MAP (Jurich et al., 2022). Three biological replicates were used. To eliminate background signal, the DMS reactivity of a PBS control was subtracted from the experimental samples. For comparisons of DMS mutational profiles between conditions, the fold change was calculated from the average DMS signal across the three replicates, eliminating bases that had exactly 0 signal from the analysis. To determine significance, an adjusted p-value was determined using the Bonferroni correction where a p-value of 0.05 was divided by the number of bases considered in the analysis. The Z-scores were then calculated using these corrected p-values (two tailed. The threshold lines were determined using the formula: log(average ± (Z-score x standard deviation)). Ratios of DMS reactivities that fall outside the threshold lines are considered significant. For reproducibility comparisons between replicates, the normalized DMS signal was generated using the box-plot method (Low and Weeks, 2010) where values larger than 1.5 times the interquartile range above the 75^th^ percentile are identified as outliers and temporarily removed from the dataset. The average of the top 10% of the remaining bases were used as the normalization factor/divisor for the entire dataset.

### Analysis of RNA abundance in cells and RNA enrichment in SGs using smFISH

Individual cells were cropped from deconvolved images using the freehand selection tool in Fiji (ImageJ). The fluorescence intensity and position of smFISH labeled RNAs were measured using the Airlocalize spot detection algorithm as described before (Trcek et al., 2017, Lionnet et al., 2011). To quantify RNAs present in the nucleus, cytoplasm and stress granules, the fluorescent spots detected by Airlocalize were manually segmented and sorted in Fiji.

### Western Blot Analysis

For western blot analysis of stably transfected U-2 OS cells expressing 6xHis-MCP-G3BP1, a fixed Region of Interest (ROI) was chosen to encompass a single protein band. The signal intensity of G3BP1 and 6xHis-MCP-G3BP1 was then measured and normalized by subtracting the signal intensity of the local background from an ROI above or below the bands of interest per lane. For ΔΔG3BP1/2 U-2 OS cells stably expressing 6xHis-MCP-G3BP1, the expression of G3BP1 was normalized by band intensities of GAPDH housekeeping gene as a loading control. For WT U-2 OS cells stably expressing 6xHis-MCP-G3BP1, the 6xHis-MCP-G3BP1 signal was normalized by the intensity of the endogenous G3BP1 used as loading controls. The fold expression of 6XHis-MCP-G3BP1 in stable clones was compared to expression of G3BP1 in WT U-2 OS cells.

### Quantifying RNA co-localization in the cytoplasm and SGs using PCC

Analyses were performed using and PCC co-localization ImageJ plugin (Bolte and Cordelières, 2006) as described before (Trcek et al., 2020). Cropped cells were used as the region of interest of PCC analysis.

### Validation of the MCP-G3BP1 cell line

Western blot analysis showed that MCP-G3BP1 expression in EF1αTET reached 136.7% of endogenous G3BP1 level (Fig. S4C). Expression of His-tagged MCP-G3BP1 also revealed co-localization between 6xHis-MCP-G3BP1 and TIA1 in ΔΔG3BP1/2 cells that lacked endogenous G3BP1 expression (Fig. S4D-F). These results confirmed that 6xHis-MCP-G3BP1 alone could form SGs and recruit SG components(Fischer et al., 2020), indicating that the fusion protein was functional.

## SUPPLEMENTAL FIGURE LEGENDS

**Figure S1: Implementing AMT-crosslinking to co-precipitate U3 snoRNA with pre-18S rRNA. A.** Calculation of ratiometric enrichment between interacting RNAs, similar to the 2(-ΔΔCq) method (Livak and Schmittgen, 2001). Fold enrichment of each RNA was first calculated as 2(Cq_target − Cq_GFP), where Cq_target is the Cq of a specific RNA (e.g., U3 snoRNA or pre-18S rRNA) recovered with probes against the target RNA (18S rRNA), and Cq_GFP is the Cq of the same RNA from the same input sample recovered with control GFP probes. The GFP pulldown thus serves as the background for non-specific co-precipitation. Ratiometric enrichment was then calculated by dividing the fold enrichment of the interacting RNA (e.g., U3 snoRNA) by that of the probe-targeted RNA (pre-18S rRNA). This normalizes for differences in probe pulldown efficiency between conditions. **B.** Cq values for the pre-18S rRNA and co-precipitated U3 snoRNA after pulldown with 18S rRNA or GFP biotinylated probes measured by RT-qPCR in unstressed and untreated WT U-2 OS cells (no AMT) or unstressed and stressed WT cells treated with AMT (AMT Unstressed and AMT Stressed, respectively). **C.** Log fold enrichment of pre-18S rRNA and co-precipitated U3 snoRNA levels. An average of six (no AMT Unstressed) to eight (AMT Unstressed and AMT Stressed) samples +/- STDEV is plotted, respectively (see Materials and Methods). **D.** Cq values for the pre-18S rRNA and co-precipitated U2 snRNA after pulldown with 18S rRNA or GFP biotinylated probes measured by RT-qPCR. **E.** Log fold enrichment of pre-18S rRNA and co-precipitated U2 snRNA levels. An average of three samples per condition +/- STDEV is plotted (see Materials and Methods). **F.** Ratiometric enrichment of U2 snRNA levels normalized by the pre-18S rRNA. A mean of three independent samples +/- STDEV is plotted. Welch’s ANOVA with Dunnett’s multiple comparison correction was used to determine significance. ns: not statistically significant, *: p < 0.05. **G.** Log fold expression of RNAs quantified by RT-qPCR relative to the *GAPDH* housekeeping gene in stressed and unstressed cells. For each condition, an average of three biological replicates +/- STDEV is shown. Dashed line in C and E demarcates no fold change.

**Figure S2: Characterization of intermolecular base pairing between in vitro transcribed *FLuc* and *RLuc* reporter RNAs using AMT crosslinking. A.** Proportion of AMT-crosslinkable sites (pyrimidines on opposite strands of base-paired dinucleotides) among all possible base-paired dinucleotide sites in the *FLuc* and *RLuc* CDS only reporter and the 3SL Sense:Antisense reporter. Each circle represents 1% of sites (100 circles total). **B.** Cq values for the *in vitro* transcribed “CDS only” reporter RNA fragment, with or without AMT, after pulldown with *FLuc* or GFP biotinylated probes, measured by RT-qPCR. **C.** Log fold enrichment of *FLuc* and co-precipitated *RLuc* from the CDS only reporter, with or without AMT. **D.** Cq values for in vitro transcribed 3SL Sense:Antisense reporter RNA fragments, with or without AMT, after pulldown with biotinylated *FLuc* or GFP probes, measured by RT-qPCR. **E.** Log fold enrichment of *FLuc* and co-precipitated *RLuc* from the 3SL Sense:Antisense reporter, with or without AMT.

**Figure S3: Evaluating intermolecular base pairing of the 3SL sense:antisense *FLuc* and *RLuc* reporter pair in EF1αTET and ΔΔG3BP1/2 EF1αTET U-2 OS cells. A.** Treatment of cells with sodium arsenite, arsenite + UV X-link, and arsenite, AMT and UV X-linking, followed by immunofluorescence against G3BP1(green) to demonstrate the SGs remain intact. Number of cells out of total inspected with the indicated phenotype is shown. DAPI stain demarcates the nuclei (blue). Scale bars: 10µm. **B.** Cq values of *FLuc* and co-precipitated RLuc of the 3SL Sense:Antisense reporter transfected into EF1αTET U-2 OS cells after the pulldown using FLuc or GFP biotinylated probes with or without AMT and sodium arsenite. The mean of three biological replicates +/- STDEV is shown. **C.** Log fold enrichment of *FLuc* and co-precipitated *RLuc* of the 3SL Sense:Antisense reporter transfected into EF1αTET U-2 OS cells, with or without AMT and sodium arsenite. The mean of three biological replicates +/- STDEV is shown. **D.** Ratiometric co-precipitation 3SL Sense:Antisense *FLuc* and *RLuc* reporter pair expressed in EF1αTET ΔΔG3BP1/2 U-2 OS cells. Per condition, an average of three biological replicates +/- STDEV of log transformed values is shown. Statistical significance was calculated using Welch’s ANOVA with Dunnett’s T3 multiple comparisons. ns: not significant. **E.** RT-qPCR analysis of *FLuc* and *RLuc* mRNA levels of the 3SL sense:antisense reporter pair after 36 hours of 1ug/mL doxycycline induction in stressed and unstressed ΔΔG3BP1/2 EF1αTET U-2 OS cells. The mean of three biological replicates ± STDEV is shown for log transformed values. qPCR quantification was done using absolute standard curves generated from dilutions of purified DNA amplicons. Significance was determined using Student’s T-test with Welch’s correction. ns: not significant. **F.** Cq values of *FLuc* and co-precipitated *RLuc* of the 3SL Sense:Antisense reporter transfected into EF1αTET ΔΔG3BP1/2 U-2 OS cells, after the pulldown using FLuc or GFP biotinylated probes with or without AMT and sodium arsenite. The average of three biological replicates +/- STDEV is shown. **G.** Log fold enrichment of *FLuc* and co-precipitated *RLuc* of the 3SL Sense:Antisense reporter transfected into EF1αTET ΔΔG3BP1/2 U-2 OS cells, with or without AMT and sodium arsenite. The average of three biological replicates +/- STDEV is shown.

**Figure S4: Establishing the MCP-G3BP1 EF1αTET cell line and evaluating intermolecular base pairing of the 3SL sense:antisense *FLuc* and *RLuc* in the prescence of MCP-G3BP1. A.** Images of U-2 OS cells ectopically expressing G3BP1:GFP and immunostained for the indicated SG marker. DAPI stain demarcates the nuclei (blue). Scale bars: 20µm. B. Images of stressed G3BP1:GFP expressing and ΔΔG3BP1/2 U-2 OS cells immunostained for the indicated SG marker (magenta). DAPI stain demarcates the nuclei (blue). Scale bars: 20µm. C. Western blot of the G3BP1 protein in clones of U-2 OS cells stably expressing 6xHis tagged MCP-G3BP1 from a CMV promoter ectopically to the endogenous G3BP1. Clone C8 marked with a red box was used for the experiments performed with the *FLuc*/*RLuc* reporter in Fig 3. Clone C8 and other tested clones also stably expresses the TET3G protein from an EF1α promoter for induction of the luciferase reporter D. Western blot of the G3BP1 protein expressed in clones of ΔΔG3BP1/2 U-2 OS cells stably expressing 6xHis-tagged MCP-G3BP1 from a CMV promoter. Clone 11 marked with a red box was used to examine the ability of 6xHis-MCP-G3BP1 to form SGs (see E, F). Clone C11 and other tested clones also stably expresses the TET3G protein from an EF1α promoter. E,F. ΔΔG3BP1/2 U-2 OS cells stably expressing 6xHis MCP-G3BP1 (C11 from D) immunostained against G3BP1 (E), TIA1 (F) and His tag in unstressed and stressed conditions. DAPI stain demarcates the nuclei (blue). Scale bars: 20µm. G. Cq values of *FLuc* and co-precipitated *RLuc* of the 3SL Sense:Antisense reporter transfected into EF1αTET MCP-G3BP1 U-2 OS cells after the pulldown using FLuc or GFP biotinylated probes with or without AMT and sodium arsenite. The average of three biological replicates +/- STDEV is shown. H. Log fold enrichment of *FLuc* and co-precipitated *RLuc* of the 3SL Sense:Antisense reporter transfected into EF1αTET MCP-G3BP1 U-2 OS cells, with or without AMT and sodium arsenite. The average of three biological replicates +/- STDEV is shown.

**Figure S5: Evaluating intermolecular base pairing of the *shu FLuc* and *RLuc* reporter pair in EF1αTET U-2 OS cells. A.** Schematic of the *shu* 3′ UTR RLuc and FLuc reporter pair tagged with 12x MS2 stem loops. **B.** Cq values of *FLuc* and co-precipitated *RLuc* of the *shu RLuc* and *FLuc* reporter pair transfected into EF1αTET U-2 OS cells after the pulldown using FLuc or GFP biotinylated probes with or without AMT and sodium arsenite. The average of three biological replicates +/- STDEV is shown. **C.** Log fold enrichment of *FLuc* and co-precipitated *RLuc* of the *shu RLuc* and *FLuc* reporter transfected into EF1αTET U-2 OS cells, with or without AMT and sodium arsenite. The average of three biological replicates +/- STDEV is shown. **D.** Ratiometric co-precipitation of the *shu FLuc* and *RLuc* reporter pair expressed in EF1αTET U-2 OS cells. Per condition, an average of three biological replicates +/- STDEV of log transformed values is shown. Statistical significance was calculated using Welch’s ANOVA with Dunnett’s T3 multiple comparisons. ns: not significant. **E.** RT-qPCR levels of the *shu RLuc* and *FLuc* reporter in EF1αTET cells after 36 hours of 1ug/mL doxycycline induction. The mean of three biological replicates ± STDEV is shown for log transformed values. qPCR quantification was done using absolute standard curves generated from dilutions of purified DNA amplicons. Significance for RT-qPCR was determined using Student’s T-test with Welch’s correction. *: p < 0.05, ns: not statistically significant.

**Figure S6: Evaluating cellular localization and intermolecular base pairing of the *shu FLuc* and *RLuc* reporter pair in EF1αTET MCP-G3BP1 U-2 OS cells. A.** Images of cells immunostained for G3BP1 (blue) and labeled with smFISH for *RLuc* (magenta) and *FLuc* (green) mRNAs in stressed EF1αTET or EF1αTET MCP-G3BP1 U-2 OS cells transfected with the *shu RLuc* and *FLuc* reporter pair. The antibody targeting the G3BP1 protein labeled the endogenous G3BP1 as well as the ectopically expressed 6xHis-MCP-G3BP1. DAPI stain demarcates the nuclei (white). Scale bar: 20 µm. **B.** Fraction of the smFISH signal from the *shu FLuc* and *RLuc* reporter mRNAs localized to SGs relative to total cellular signal in WT (green) and 6xHis-MCP-G3BP1-expressing (magenta) cells. A total of 56 EF1αTET and 57 EF1αTET MCP-G3BP1 cells were analyzed. **C.** Cq values of *FLuc* and co-precipitated *RLuc* of the *shu* reporter transfected into MCP-G3BP1 EF1αTET U-2 OS cells, after the pulldown using *FLuc* or GFP biotinylated probes with or without AMT and sodium arsenite. The average of three biological replicates +/- STDEV is shown. **D.** Log fold enrichment of *FLuc* and co-precipitated *RLuc* of the *shu* reporter transfected into MCP-G3BP1 EF1αTET U-2 OS cells, with or without AMT and sodium arsenite. The average of three biological replicates +/- STDEV is shown. **E.** Ratiometric enrichment of *shu FLuc* and *RLuc* reporter pair expressed in MCP-G3BP1 EF1αTET U-2 OS cells. Per condition, an average of three biological replicates +/- STDEV of log transformed values is shown. Statistical significance was calculated using Welch’s ANOVA with Dunnett’s T3 multiple comparisons correction. **F.** RT-qPCR levels of the *shu RLuc* and *FLuc* reporter in MCP-G3BP1 EF1αTET cells. The mean of three biological replicates ± STDEV is shown for log transformed values. qPCR quantification was done using absolute standard curves generated from dilutions of purified DNA amplicons. Significance for RT-qPCR was determined using Student’s T-test with Welch’s correction. *: p < 0.05, ns: not statistically significant.

**Figure S7: Evaluating intermolecular base pairing between *AHNAK* and *NORAD* in WT and ΔΔG3BP1/2 U-2 OS cells. A.** Minimal interaction energy prediction of intermolecular RNA:RNA interactions with intaRNA (Mann et al., 2017). The interaction energy in kcal/mol is the minimal free energy of RNA base pairing interactions for an RNA pair at intersecting regions of the two RNAs. A lower interaction energy suggests a more energetically favorable base pairing interaction. **B, C.** Proportion of AMT-crosslinkable sites (pyrimidines on opposite strands of base-paired dinucleotides) among all possible base-paired dinucleotide sites in *AHNAK* (B) and *NORAD* (C). Each circle represents 1% of sites (100 circles total). **D.** Schematic depicting co-precipitation of lnc *NORAD* (green) with *AHNAK* mRNA (magenta) hybridized with biotinylated oligonucleotides (oligos with yellow circles). Co-precipitated RNAs were reverse transcribed and their levels examined using RT-qPCR analysis (blue polymerase and blue amplified DNA amplicons). **E.** Absolute quantification of *NORAD* RT-qPCR amplicons per ng of input RNA in WT and ΔΔG3BP1/2 U-2 OS cells. Sample Cq values were matched with Cq values from a standard curve of the same purified DNA amplicon. The mean of four replicates ± STDEV per amplicon is shown. **F.** Ratiometric enrichment of *NORAD* relative to *AHNAK* levels in WT U-2 OS cells with or without AMT and sodium arsenite. For each condition, an average of three biological replicates +/- STDEV is shown where possible. Amplicons without 3 replicates indicate replicate samples that did not have detectable Cq values for analysis. Statistical significance was evaluated using Welch’s ANOVA test with Dunnett’s T3 multiple comparisons correction. **G.** Log fold enrichment for the *AHNAK* mRNA and co-precipitated *NORAD* lncRNA in WT cells with or without AMT and sodium arsenite. **H.** Ratiometric enrichment of *NORAD* relative to *AHNAK* in ΔΔG3BP1/2 U-2 OS cells with or without AMT and sodium arsenite. For each condition, an average of three biological replicates +/- STDEV is shown where possible. Amplicons without 3 replicates indicate replicate samples that did not have detectable Cq values for analysis. Statistical significance was evaluated using Welch’s ANOVA test with Dunnett’s T3 multiple comparisons correction. **I.** Log fold enrichment for the *AHNAK* mRNA and co-precipitated *NORAD* lncRNA in ΔΔG3BP1/2 U-2 OS cells with or without AMT and sodium arsenite. *: p < 0.05, **: p < 0.01.

**Figure S8: Evaluating colocalization between *AHNAK* and *TLN1* and *DYNC1H1* in WT and ΔΔG3BP1/2 U-2 OS cells. A-D.** Images of cells immunostained for G3BP1 (blue) and labeled with smFISH for *AHNAK* (green) and *TLN1* or *DYNC1H1* (magenta) in unstressed (A,C) and stressed (B,D) WT U-2 OS cells. DAPI stain demarcates the nuclei (white). Zoomed-in images (white squares) show *AHNAK*, *TLN1* or *DYNC1H1* mRNAs dispersed in the cytosol or in co-localized SGs. Scale bars: 1 and 10 µm in small and big images, respectively. **E, F.** Quantification of *TLN1* (E) and *DYNC1H1* (F) in different cellular compartments in stressed WT U-OS cells. 20 cells were analyzed per condition. **G-J** Images of cells immunostained for G3BP1 (blue) and labeled with smFISH for *AHNAK* (green) and *TLN1* or *DYNC1H1* (magenta) in unstressed (G, I) and stressed (H, J) ΔΔG3BP1/2 U-2 OS cells. DAPI stain demarcates the nuclei (white). Zoomed-in images (white squares) show *AHNAK*, *TLN1* or *DYNC1H1* mRNAs dispersed in the cytosol. Scale bars: 1 and 10 µm in small and big images, respectively. **K,L.** Co-localization of *AHNAK* with *TLN1* (G) and *AHNAK* with *DYNC1H1* (J) in the cytoplasm of unstressed and stressed WT and ΔΔG3BP1/2 U-2 OS cells. SG ROIs were cropped out to determine co-localization between *AHNAK* and *TLN1* or *AHNAK* and *DYNC1H1* specifically in SGs for stressed WT cells. For each condition, a minimum of 63 unstressed or stressed cells were analyzed. The dashed line indicates the average PCC value of doubly stained *AHNAK* smFISH in WT U-2 OS cells (from Fig. 4B,C). Statistical significance was examined using Welch’s ANOVA with Dunnett’s T3 multiple comparisons correction. ****: p < 0.0001. *: p < 0.05. **M**. Relative RT-qPCR levels of *AHNAK*, *NORAD*, *DYNC1H1* and *TLN1* in unstressed and stressed WT and ΔΔG3BP1/2 U-2 OS cells, relative to *GAPDH*. For each condition, an average of three biological replicates +/- STDEV is shown.

**Figure S9: Comparison of DMS reactivity of the *AHNAK* 3′UTR between biological replicates in unstressed and stressed WT and ΔΔG3BP1/2 cells. A,B.** Correlation of normalized DMS reactivity across biological replicates of the *AHNAK* 3′UTR in unstressed (U1,2,3) and stressed (S1,2,3) WT (A) and ΔΔG3BP1/2 (B) U-2 OS cells. The dark blue line represents the linear regression of all bases across replicate pairs. The correlation coefficient (r) is shown in red.

**Figure S10: Comparison of DMS reactivity of *NORAD* ND3 and ND4 between biological replicates in unstressed and stressed WT and ΔΔG3BP1/2 cells. A,B.** Correlation of normalized DMS reactivity across biological replicates of *NORAD* ND3 and ND4 in unstressed (U1,2,3) and stressed (S1,2,3) WT (A) and ΔΔG3BP1/2 (B) U-2 OS cells. The dark blue line represents the linear regression of all bases across replicate pairs. The correlation coefficient (r) is shown in red.

## SUPPLEMENTAL TABLES

**Table S1.** Biotinylated probes for AMT crosslinking pulldown

**Table S2.** RT-qPCR and in-vitro transcription primers

**Table S3.** smFISH probe sequences

**Table S4.** Targeted DMS-MaPseq primers

**Table S5.** rRNA subtraction oligos (from (Adiconis et al., 2013))

## Notes

### Competing Interest Statement

The authors have declared no competing interest.

### Summary of Updates

We have provided new data that expand the controls for the AMT crosslinking-dependent characterization of intermolecular base-pairing interactions, both in vitro and in cells

